# Single-cell transcriptomic analysis of pea shoot development and cell-type-specific responses to boron deficiency

**DOI:** 10.1101/2023.06.14.545017

**Authors:** Xi Chen, Yanqi Ru, Hirokazu Takahashi, Mikio Nakazono, Sergey Shabala, Steven M. Smith, Min Yu

**Author notes:** Correspondence: Steven M. Smith; Min Yu. E-mail address (X.C.); (Y.R.);<u>a-</u> <u>u.ac.jp</u> (H.T.); (M.N.); (S.S.); (S.M.S.); (M.Y.).

## Abstract

Understanding how nutrient stress impacts plant growth is fundamentally important to the development of approaches to improve crop production under nutrient limitation. Here we applied single-cell RNA sequencing to shoot apices of *Pisum sativum* grown under boron (B) deficiency. We identified up to fifteen cell clusters based on the clustering of gene expression profiles and verified cell identity with cell-type-specific marker gene expression. Different cell types responded differently to B deficiency. Specifically, the expression of photosynthetic genes in mesophyll cells (MCs) was down-regulated by B deficiency, consistent with impaired photosynthetic rate. Furthermore, the down-regulation of stomatal development genes in guard cells (GCs), including homologs of *MUTE* and *TOO MANY MOUTHS*, correlated with a decrease in stomatal density under B deficiency. We also constructed the developmental trajectory of the shoot apical meristem (SAM) cells and a transcription factor (TF) interaction network. The developmental progression of SAM to MC was characterized by up-regulation of genes encoding histones and chromatin assembly and remodeling proteins including homologs of FASCIATA1 (FAS1) and SWITCH DEFECTIVE/SUCROSE NON-FERMENTABLE (SWI/SNF) complex. However, B deficiency suppressed their expression, which helps to explain impaired SAM development under B deficiency. These results represent a major advance over bulk-tissue RNA-seq analysis in which cell-type-specific responses are lost and hence important physiological responses to B deficiency are missed. The reported approach and resources have potential applications well beyond *P. sativum* species and could be applied to various legumes to improve their adaptability to multiple nutrient or abiotic stresses.

## Introduction

Boron (B) deficiency is a common problem causing reduced arable crop yields (Goldbach and Wimmer, 2007). Although B has been identified as an essential element for plants for 100 years (Warington, 1923), the fine-print of its operation in plants at the molecular level remains to be revealed, and any putative B receptors remain to be discovered. These challenges are mostly due to the fact that B is involved in numerous physiological and biochemical responses, making the separation of direct and indirect effects difficult. One of the direct pieces of evidence for the essentiality of B in plants is its role in maintaining the structural stability of cell walls and membranes (O’Neill et al., 2001; Voxeur and Fry, 2014). In plants, 90% of B is present in the cell wall, where it covalently cross-links two rhamnogalacturonan II (RG-II) monomers to form the dimeric RG-II-B, supporting cell wall structure and function (O’Neill et al., 1996). This role makes B essential for controlling plant architecture. However, whether different plant tissues exhibit different responses to B deficiency stress, whether different cell types have varying tolerances to B deficiency, and how aerial tissues adopt different strategies to adapt to B stress remains to be further investigated.

In higher plants, the maturation of the shoot apical meristem (SAM) involves two main processes: maintaining the activity of the pluripotent stem cell population and the continuous differentiation of the meristem for the reiterative formation of lateral organs (Bowman and Eshed, 2000). The internal regulatory mechanisms of the SAM typically involve the activation of the CLAVATA3/EMBRYO SURROUNDING REGION (CLE) family by WUSCHEL (WUS) in the organizing center (OC), maintaining stem cells through a signaling network. External regulation of the SAM includes the effects of environmental signals acting to influence the distribution of endogenous plant hormones, such as auxins and cytokinins, as well as modulation of signal perception to activate *WUS* expression (Bowman and Eshed, 2000; Zhang et al., 2017). Another homeodomain transcription factor (TF) isolated from a *Zea mays* mutant, KNOTTED1 (KN1) (*Arabidopsis thaliana* ortholog, SHOOTMERISTEMLESS/STM), plays a role in preventing premature differentiation of meristem cells (Carles and Fletcher, 2003). Research has shown that meristem development is particularly sensitive to B deficiency (Poza-Viejo et al., 2018). The reduced-B-uptake mutant *tassel-less1* (*tls1*) in *Z. mays* exhibits defective SAM structure due to transcriptional repression of the meristem maintenance gene *KNOTTED1* (*KN1*) and cell cycle genes (Matthes et al., 2022). However, how B deficiency affects the regulation of meristem development and the key TFs involved in this process have yet to be discovered.

Single-cell RNA-sequencing (scRNA-seq) achieves single-cell resolution for transcriptome sequencing of tissues. In addition to capturing even rare cell types, the spatiotemporal resolution of scRNA-seq helps us understand the relationships between cell types in space and time. By constructing developmental trajectories of cell types, we can reconstruct the differentiation process of a given tissue and the potential regulatory genes involved (Shaw et al., 2020). Currently, several studies have applied scRNA-seq or single-nucleus RNA-seq (snRNA-seq) to explore differences in cell types and developmental and differentiation trajectories in various plant species and tissues, including *Arabidopsis thaliana* shoots and roots (Denyer et al., 2019; Zhang et al., 2021b), maize (*Z. mays*) shoots (Satterlee et al., 2020), rice (*Oryza sativa*) shoots and roots (Liu et al., 2021b; Wang et al., 2021), poplar (*Populus*) stems and shoots (Chen et al., 2021; Conde et al., 2022), peanut (*Arachis hypogaea*) leaves (Liu et al., 2021a), tea (*Camellia sinensis*) leaves (Wang et al., 2022), *Medicago truncatula* root nodules (Ye et al., 2022), soybean (*Glycine max*) nodules (Liu et al., 2023) and wheat (*Triticum aestivum*) roots (Zhang et al., 2023). A few studies have also focused on the heterogeneous response of plant tissues under stress which include *Z. mays* roots in response to nitrate (Li et al., 2022), strawberry (*Fragaria vesca*) leaves in response to *Botrytis cinerea* infection (Bai et al., 2022), Chinese cabbage (*Brassica rapa*) leaves in response to heat stress (Sun et al., 2022a), rubber tree (*Hevea brasiliensis*) leaves in response to powdery mildew infection (Liang et al., 2023). However, the application of single cell sequencing technology in the response of shoots to nutrient stress needs further investigation. scRNA-seq is rapidly advancing in the application of plant genomics research and is becoming an important tool for studying cell- and tissue-specific gene functions in plants.

Cell- and tissue-specific marker genes have been widely reported in well-annotated species such as *A. thaliana*, *O. sativa* and *Z. mays*. They have been identified and validated by several techniques including reporter gene expression in transgenic plants, *in situ* hybridization to tissue sections, and laser capture microdissection (LCM) (Berkowitz et al., 2021; Bezrutczyk et al., 2021; Gala et al., 2021; Zhang et al., 2021a; Zhang et al., 2021b). However, reports of marker genes in legume plants, particularly in aerial tissues, are scarce. scRNA-seq enables the identification of new markers in other less-extensively studied species such as *P. sativum* by analyzing the transcriptomes of different cell types. LCM allows us to easily obtain different tissues within the shoot apex to verify identification of new marker genes in such plants (Takahashi et al., 2010). With this powerful technology platform, it is now possible to study tissue-specific responses of agronomically-important plants to biotic and abiotic stresses and uncover differential gene expression responses in specific cell types.

As the subject of Mendel’s genetic research, pea (*Pisum sativum*) has become one of the more important subjects for plant genetics and developmental research due to its nitrogen-fixing ability, high nutritional value and agricultural importance (Wong et al., 2008), with the world production of commercial *P. sativum* species estimated at 36.91 million tons (MT) in 2017 (Tassoni et al., 2020). The genomes of different *P. sativum* varieties have been sequenced, and the information is constantly being refined (Kreplak et al., 2019; Yang et al., 2022). However, due to the large size of the *P. sativum* genome, the high proportion of repetitive sequences, and difficulties in applying functional genomics approaches, the annotation of *P. sativum* genes is lacking relative to soybean (*G. max*), *M. truncatula* and *Lotus japonicus* (Smýkal et al., 2012). Therefore, the introduction of new omics technologies can help further develop *P. sativum* genomic resources and provide more information for genetic improvement and germplasm utilization.

In this study, we used a single-cell transcriptome approach to characterize the responses of shoot apical cells of *P. sativum* to B availability. The results reveal critical genes and regulatory factors related to shoot development in response to B deficiency and provide insights into strategies by which plants adapt to B deficiency thus offering breeders a set of specific targets for genetic improvement. The approach and resources could be applied to various legumes to improve their adaptability to multiple nutrient or abiotic stresses.

## Results

### A single-cell atlas of *P. sativum* shoot apices

Pea (*P. sativum*) seeds were germinated in 0.5 mM CaCl_2_ solution for 2 d, after which the seedlings were transferred to hydroponic culture in a 1/4 strength modified Hoagland nutrient solution with either 25 μM B (B25) or no B added (B0) for an additional 10 d. More than 50 shoot apices were collected for each treatment. The apices were chopped with a razor blade and quickly transferred to 5 ml of cellulose and pectinase enzyme solution. After incubating for 1 h, cell number and viability were determined using light microscopy (Figure S1a) and staining with Trypan blue, respectively. More than 20,000 cells from each sample were labeled using the 10x Genomics scRNA-seq platform. Following the construction of cDNA libraries and Illumina high-throughput sequencing, gene expression levels were assessed by analyzing Unique Molecular Identifier (UMI) counts. The median number of genes detected per cell was 3,416 (B0) and 2,963 (B25), and the median UMI counts per cell were 10,567 (B0) and 8,822 (B25). After filtering, a total of 14,493 high-quality cells were obtained from B25, with 25,390 genes mapped, while 9,212 high-quality cells were obtained from B0, with 25,418 genes (Table S1). Seventy-seven percent of the genes identified by scRNA-seq were also detected in previous analysis of bulk RNA-seq data (Figure S1b) (Chen et al., 2022). Additionally, Pearson correlation analysis was performed on the 4,000 genes showing the greatest fold change (FC) in response to B deficiency in bulk RNA-seq and aggregated scRNA-seq data, revealing a correlation coefficient of 0.6 between the two platforms. Our results demonstrate a high degree of correlation between the genes induced by B deficiency across the two platforms.

The transcriptome data were then subjected to Principal Component Analysis (PCA) dimensionality reduction and clustering analysis, resulting in 15 distinct cell clusters, which were observed in both B25 and B0 (Figure S2a-b). To further elucidate the similarities and differences among cell clusters, we used the Uniform Manifold Approximation and Projection (UMAP) to visualize the cell clusters (Figure 1a). We used homologous genes in *P. sativum* of previously published marker genes from *A. thaliana* or *Z. mays* to identify each cell cluster (Figure 1b, Table S2). In MCs, *CHLOROPHYLL A-B BINDING PROTEIN CP29.1* (*Psat7g245920*, *PsLHCB4.1*), *CHLOROPHYLL A-B BINDING PROTEIN 215* (*Psat2g004560*, *PsCAB215*), *PHOTOSYSTEM I CHLOROPHYLL A/B-BINDING PROTEIN 2* (*Psat1g097880*, *PsLHCA2*), and *PHOTOSYSTEM I REACTION CENTER SUBUNIT III* (*Psat1g169240*, *PsPSAF*) were highly expressed. In epidermal cells (ECs), *HYDROXYSTEROID DEHYDROGENASE 1* (*Psat7g037080*, *PsHSD1*), *LONG CHAIN ACYL-COA SYNTHETASE 1* (*Psat6g160800*, *PsLACS1*), and *PALMITOYL-PROTEIN THIOESTERASE-DOLICHYL PYROPHOSPHATE PHOSPHATASE FUSION 1* (*Psat0s4925g0080*, *PsPDF1*) were highly expressed. In guard cells (GCs), *EPIDERMAL PATTERNING FACTOR 1* (*Psat5g271480*, *PsEPF1*), *TRANSCRIPTION FACTOR MUTE* (*Psat1g180920*, *PsMUTE*), and *TRANSCRIPTION FACTOR FAMA* (*Psat0s3505g0040*, *PsFAMA*) were highly expressed. In vascular cells (VCs), *HOMEOBOX-LEUCINE ZIPPER PROTEIN ATHB-8* (*Psat5g020800*, *PsATHB-8*), *LEUCINE-RICH REPEAT RECEPTOR-LIKE PROTEIN KINASE TDR* (*Psat6g142440*, *PsTDR*), *COPPER TRANSPORT PROTEIN CCH* (*Psat6g093320*, *PsCCH*), *AUXIN-RESPONSIVE PROTEIN IAA27* (*Psat7g014720*, *PsIAA27*), and *PROTEIN SHORT-ROOT* (*Psat0s2268g0040*, *PsSHR*) were highly expressed. *GIBBERELLIN 2-BETA-DIOXYGENASE 1* (*Psat1g189360*, *PsGA2OX1*), *HOMEOBOX PROTEIN KNOTTED-1-LIKE 3* (*Psat6g028400*, *PsKNAT3*), *GLUTATHIONE S-TRANSFERASE U8* (*Psat6g170320*, *PsGSTU8*), and *9-CIS-EPOXYCAROTENOID DIOXYGENASE NCED1* (*Psat1g001480*, *PsNCED1*) were used as markers for SAM (Figure 1b, Table S2). Furthermore, significant expression of marker genes associated with cell proliferation was noted. *HIGH MOBILITY GROUP B PROTEIN 6* (*Psat7g026240*, *PsHMGB6*), *INFLORESCENCE MERISTEM RECEPTOR-LIKE KINASE 2* (*Psat7g249120*, *PsIMK2*), and *UBIQUITIN-CONJUGATING ENZYME E2 20* (*Psat7g179720*, *PsUBC20*) serve as marker genes for proliferating cells (PCs), with notable expression in clusters 9 and 11 (Figure 1b, Table S2), indicating that these two cell clusters are engaged in cell division.

**Figure 1.**
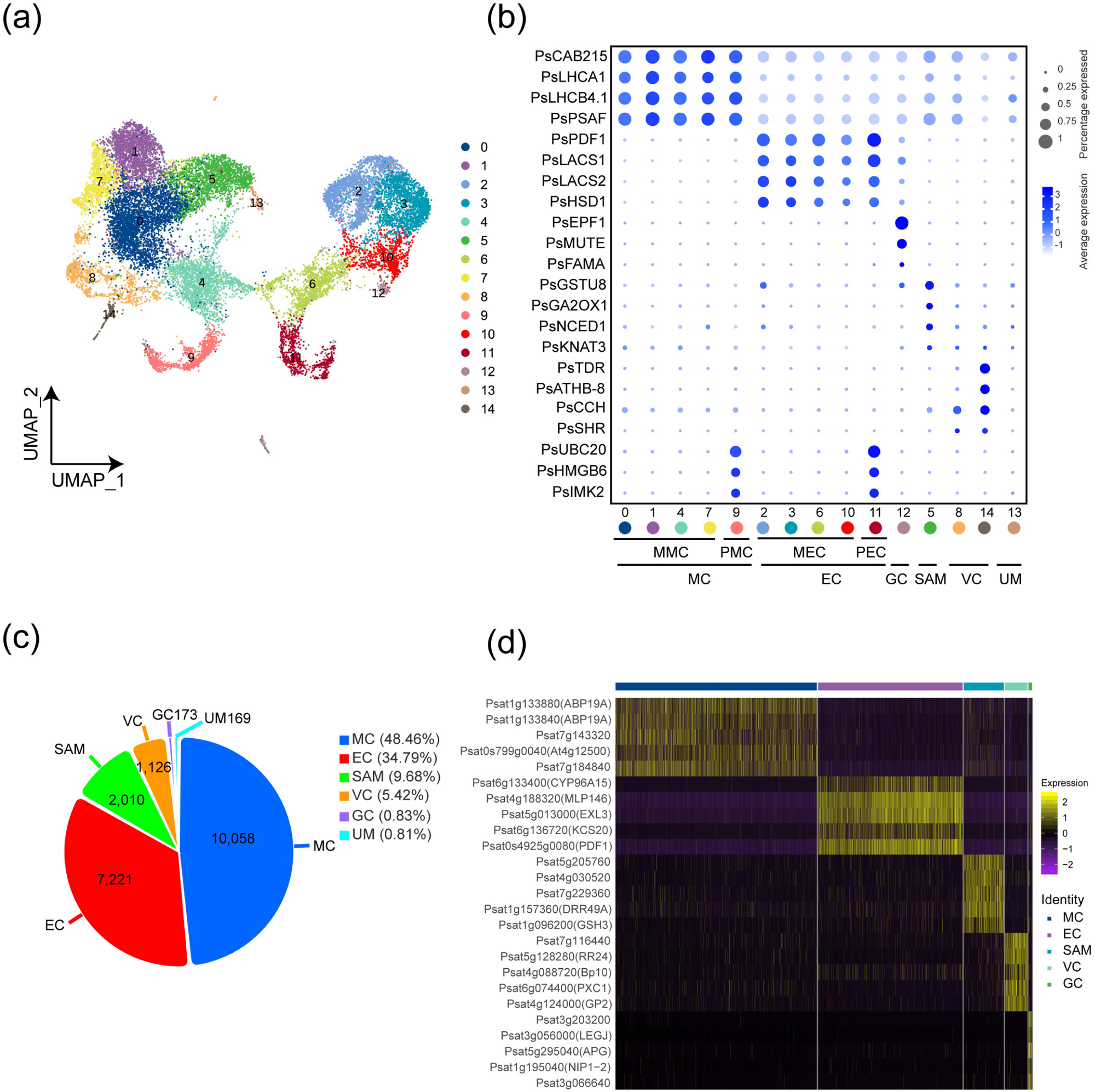
Classification of single-cell transcriptomes and cellular heterogeneity in pea (*P. sativum*) shoot apices. (a) UMAP visualization of 15 cell clusters from 20,757 cells from B25 and B0. Each dot represents an individual cell. (b) Expression pattern of cell-type-specific marker genes in each cell cluster. The dot size illustrates the percentage of cells expressing the gene, while the color intensity indicates the average expression level. Information on marker genes is given in Table S2. (c) Pie chart showing cell types, the numbers of cells, and their proportions. (d) Heatmap showing five characteristically up-regulated DEGs for all cells in each of five known cell types. Gene expression levels have been normalized by Z-score. MC, mesophyll cell; MMC, mature mesophyll cell; PMC, proliferating mesophyll cell; EC, epidermal cell; MEC, mature epidermal cell; PEC, proliferating epidermal cell; GC, guard cell; SAM, shoot apical meristem; VC, vascular cell; UM, unknown meristem.

In summary, we identified clusters 0, 1, 4, 7, and 9 as MC (10,058 cells), clusters 2, 3, 6, 10, and 11 as EC (7,221 cells), with 9 and 11 being PC of MC and EC, respectively. Cluster 12 was identified as GC (173 cells), cluster 5 as SAM (2,010 cells), and clusters 8 and 14 as VC (1,126 cells) (Figure 1c). Cluster 13 (169 cells) could not be successfully identified due to insufficient marker genes. However, it was classified as an unknown meristem (UM) due to its transcript similarity to SAM cells (Figure 1a, c).

We used LCM to capture SAM, MC, and vascular bundle tissues from apices of *P. sativum* (Figure 2a-c), and the expression of the marker genes was validated by quantitative real-time PCR (qRT-PCR). We found that *PsKNAT3*, a gene that encodes a member of the KNOX (KNOTTED1-like homeobox) family that maintains stem cell stability, as well as its downstream hormone signaling regulator *PsGA2OX1* and *PsNCED1*, regulated by CUC2 (CUP-SHAPED COTYLEDON 2) and BRC1 (BRANCHED1), were highly expressed in the SAM (Figure 2d). *PsLHCB4.1*, *PsLHCA1*, and *PsCAB215*, functioning in photosynthesis, were highly expressed in MC (Figure 2e), while *PsSHR*, *PsATHB-8*, and *PsTDR* (*TDR/PXY*, *TDR/PHLOEM INTERCALATED WITH XYLEM*), which are

**Figure 2.**
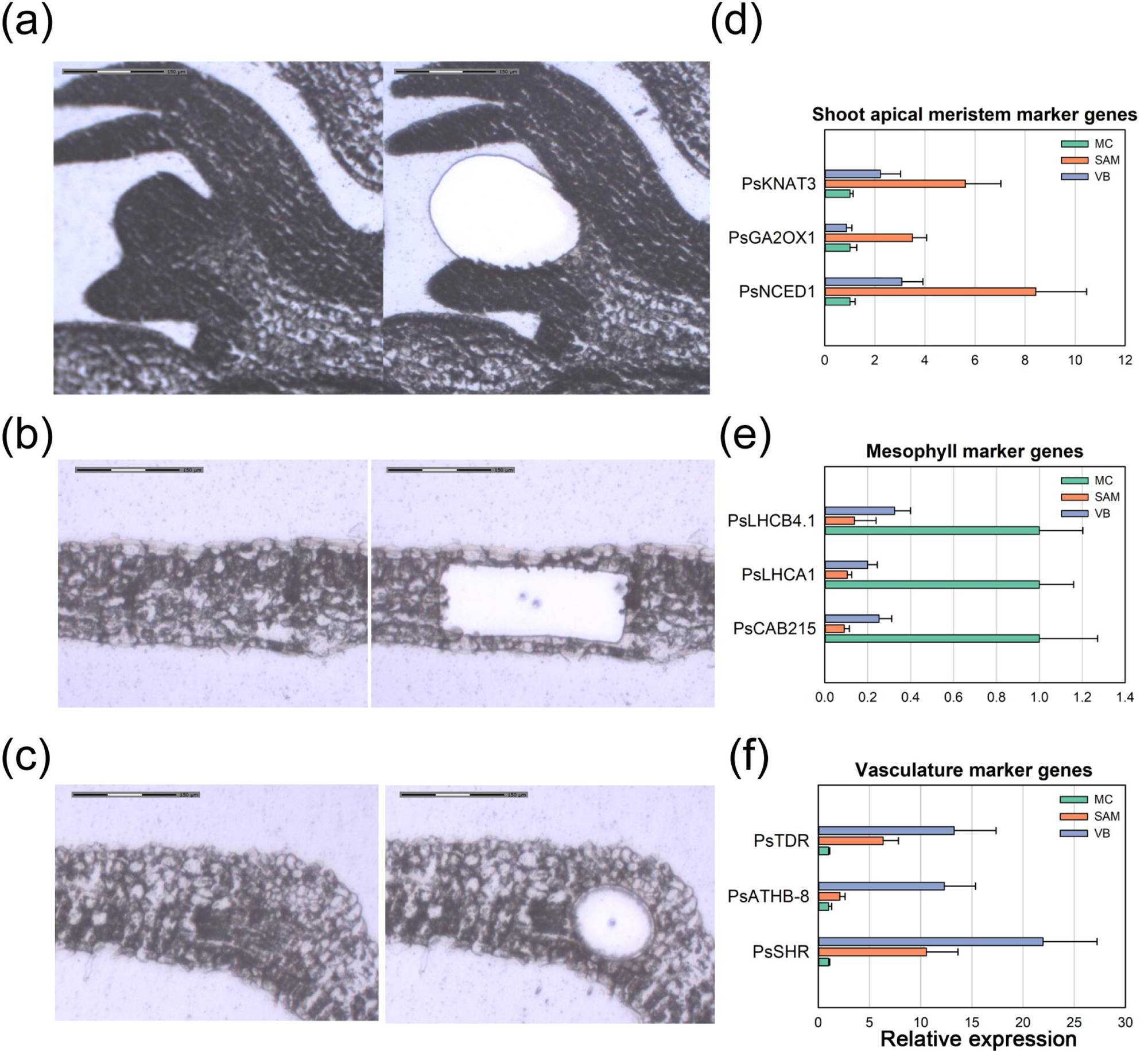
LCM of shoot apex and RT-qPCR showing the relative expression of marker genes in dissected tissues. (a-c) Shoot apical meristem (SAM, a), mesophyll cell (MC, b) and vascular bundle (VB, c) before and after LCM. (d-f) Relative expression of marker genes for SAM (d), MC (e) and VB (f). Primer sequences are given in Table S11. Actin gene in pea was used as a reference. Results were normalized relative to the expression in MC. Values represent mean ± SD (n = 3). Scale bars (a-c): 150 μm. involved in plant vascular development and differentiation, were highly expressed in the vascular bundle (Figure 2f). These results validate the reliability of our cell classification and demonstrate the accuracy of the new marker genes for cell-type-specific expression identified from this study (Figure 3). Our identification of cell types in *P. sativum* using scRNA-seq, as well as the validation of specific marker genes for these cell types, will be beneficial for recognizing different cell types in *P. sativum* and other legume plants in the future, as well as for further analyzing transcript level differences between tissues. Functional genomics in legumes is difficult and relatively little research has been conducted so the identification of cell-type-specific marker genes will greatly facilitate research in *P. sativum* and other leguminous plants.

**Figure 3.**
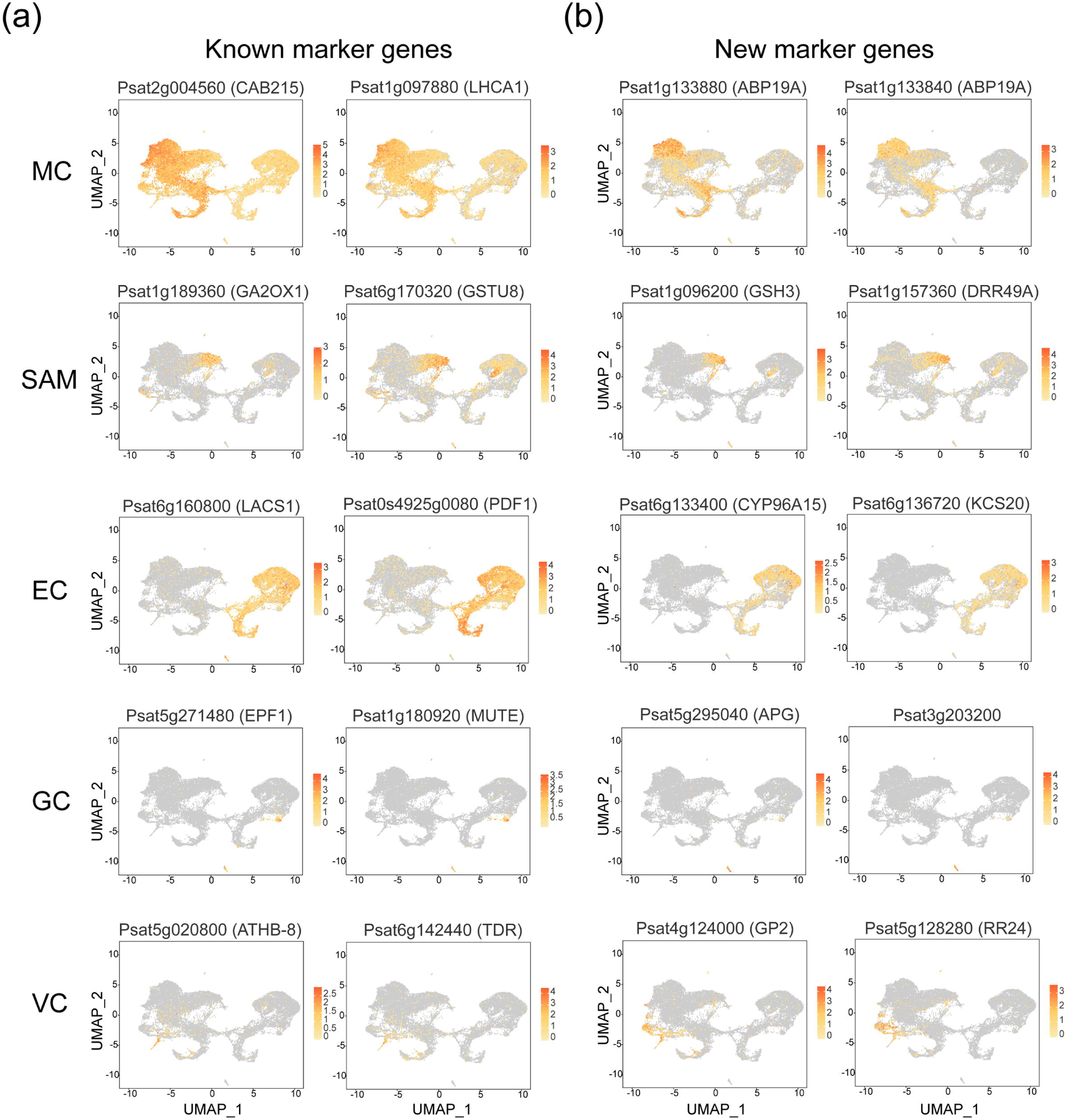
Expression of known marker genes (a) and new marker genes (b). UMAP plot illustrating the spatial distribution of expression of selected marker genes among different cell types. Color intensity reflects the level of gene expression. Gene information is given in Table S4.

To further verify the accuracy of cell type identification and investigate the molecular characterization of each cell type, we applied the Wilcoxon Rank Sum test in Seurat (R package) to analyze differentially expressed gene (DEG) expression among different cell types. We then screened for up-regulated genes in one type compared to others (these genes must be expressed in 25% of cells in the target or control type, with a *P*-value ≤ 0.01, and a log_2_FC ≥ 0.36) and showed the 5 most highly up-regulated DEGs in the five known cell types (Figure 1d). Subsequently, we conducted Gene Ontology (GO) and Kyoto Encyclopedia of Genes and Genomes (KEGG) enrichment on the up-regulated cell-type-specific genes (Figure S3, Figure S4). The potential biological functions revealed through the enriched gene pathways helps us to determine the accuracy of cell type identification. In MC, up-regulated genes were specifically enriched in photosynthesis, amino acids, and carbon metabolism, indicating that these genes are also related to the primary functions such as energy metabolism. In SAM, up-regulated genes were specifically enriched in pathways such as plant hormone signal transduction and linolenic acid metabolism, revealing the key role of hormone signals in the shoot apex. In VC, genes involved in generating proton motive force via V-type H^+^-transporting ATPase were enriched, potentially providing energy for the uptake of inorganic and organic solutes. In EC, fatty acid metabolism pathways were enriched consistent with a crucial role in cuticle biosynthesis (Figure S3, Figure S4).

### Responses to B deficiency in a cell-type-specific manner

By comparing the UMAP results of B25 and B0, we found that both include 15 cell clusters, indicating that the cell classification was not affected by B deficiency (Figure 4a, f, g). However, the transcript profiles of different cell types in response to B deficiency revealed some potentially important differences. The total number of captured cells in the B-deficient samples was lower, and the cell proportion also changed after B deficiency (Figure S2a). Boron deficiency resulted in a higher proportion of SAM and MC, while the proportion of EC and VC decreased (Table S3). Since the gene expression fluctuation in each cell is more sensitive in scRNA-seq compared to bulk RNA-seq, we used a lower threshold when screening for DEGs to avoid missing some potentially important DEGs. Genes with |log_2_FC| ≥ 0.36 and *P* < 0.05 were considered as DEGs. We identified the following numbers of DEGs in different cell types: 3,402 in MC (2,593 up, 809 down); 3,046 in VC (1,872 up, 1,174 down); 2,640 in EC (1,993 up, 647 down); 2,246 in GC (1,380 up, 866 down); 2,230 in SAM (1,407 up, 823 down); 620 in UM (158 up, 462 down) (Figure 4b, Table S4).

**Figure 4.**
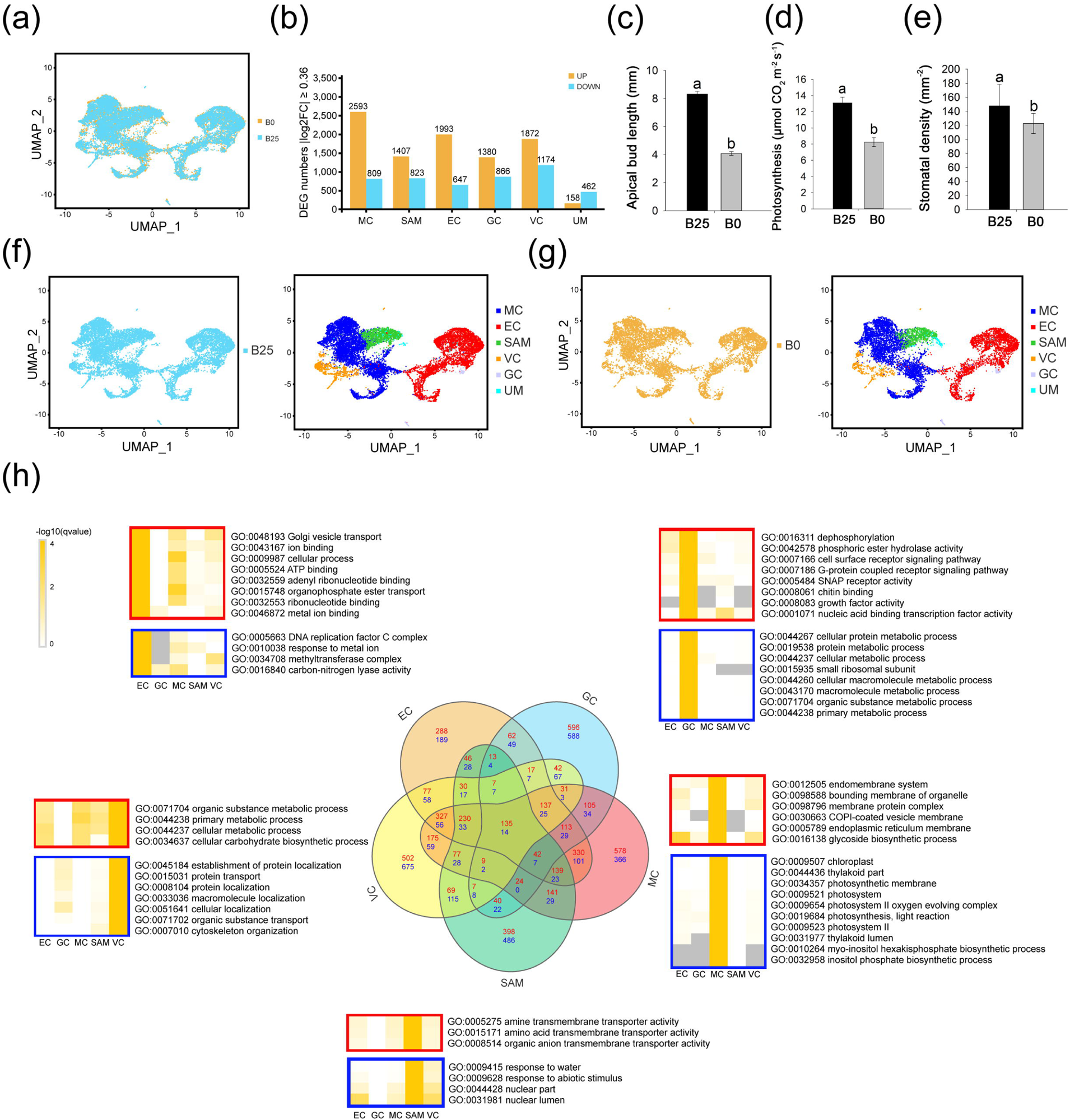
Cell-type-specific responses to B deficiency. (a) UMAP plot of pea shoot apex cells of B25 (blue) treatment overlaid upon and B0 (orange) treatment (B25, 12 618 cells; B0, 8 139 cells). (b) Numbers of DEGs (up, orange; down, blue) between B25 and B0 in six cell types. The detailed information of all the DEGs is given in Table S4-6. (c) Apical bud (n = 12) length difference between B25 and B0 treatments. (d) Photosynthesis rate (n = 6) difference in 18-day leaves from B25 and B0 treatments. (e) Stomatal density (n = 24) difference in 18-day treatments. Values represent mean ± SD. Lower-case letters indicate significant differences: Duncan’s test (P < 0.05). (f-g) UMAP visualization of individual cells and distribution of identified cell clusters (color coded) in B25 (f) and B0 (g) samples. (h) Venn diagram showing numbers of overlapping B-responsive DEGs (red, up in B0; blue, down in B0) between five known cell types (center) and GO enrichment analysis (FDR ≤ 0.05) for B-responsive DEGs for each cell-type (red boxes, up in B0; blue boxes, down in B0).

### Cell wall genes in different cell types

Since the most significant role of B is in maintaining cell wall structure (O’Neill et al., 2001), we analyzed the transcript levels of cell wall-related genes in different cell types under B deficiency. Among them, cellulose synthase genes (*CesA*) were up-regulated in all cell types, while some pectin synthesis-related genes (*GAUTs*, *GALACTURONOSYLTRANSFERASE*) were up-regulated in MC, VC, and SAM. In contrast, some pectin synthesis-related genes (*PMEs*) were down-regulated in SAM and GC. Cellulose degradation or modification genes (*XTHs*) were up-regulated in EC and GC. Polygalacturonase-related genes (*GPs*) were up-regulated in all cell types. However, cell wall loosening *EXPANSIN* genes were down-regulated in all cell types (Table S8). Boron deficiency led to the remodeling of cell walls in different cell types of the shoot apex (Table S8, Chen et al., 2022). This deficiency also caused changes in pectin content and the pectic polysaccharide network, which in turn altered properties and compromised the stability of the cell wall (Fleischer et al., 1999). Boron deficiency could significantly influence cell wall integrity and function. The extent and nature of these effects, however, appear to be cell-type-specific, underscoring the complexity of B’s role in cell wall maintenance and development.

### Analysis of DEGs in different cell types

We conducted GO enrichment analyses on these DEGs and showed specific enrichment in GO terms across different cell types (Figure 4h). Boron deficiency induced up-regulation of genes related to Golgi vesicle transport, ion binding, and ATP binding functions in EC, but led to down-regulation of genes related to DNA replication and nucleotide excision repair. In EC, *PsPDF1*, which is related to epidermal-specific features and regulates epidermal development, was down-regulated by B deficiency (Table S4).

In VC, *SHORT ROOT* (*SHR*) genes (*Psat0s2268g0040*, *PsSHRa*; *Psat0s280g0040*, *PsSHRb*) regulating vascular tissue development were suppressed by B deficiency (Table S4), highlighting the importance of B in vasculature development. Boron transporter genes *PsBOR2* and *PsBOR6* were up-regulated in MC, SAM, EC, and VC under B deficiency (Table S7). This is consistent with previous results obtained from bulk RNA-seq (Chen et al., 2022). Plasma membrane intrinsic protein genes (*PIP*s) were also up-regulated in VC which indicates the regulation of water balance under B deficiency (Table S7).

In GC, B deficiency up-regulated genes involved in dephosphorylation, cell surface receptor signaling pathways, and growth factor activity. The up-regulation of these genes may help activate the nutrient-stress adaptation mechanisms in GC, enhance signal transduction to adapt to stress, while the enrichment results of down-regulated genes in GC indicate that protein synthesis and degradation processes, as well as amino acid synthesis pathways, are depressed in GC (Figure 4h, Table S5-6). The *TRANSCRIPTION FACTOR SPEECHLESS* (*Psat6g038600*, *PsSPCH*), related to stomatal development, was up-regulated in GC, while *TOO MANY MOUTHS* (*Psat5g302680*, *PsTMM*) and *TRANSCRIPTION FACTOR MUTE* (*Psat1g180920*, *PsMUTE*) were down-regulated in GC (Table S4). Boron deficiency may suppress stomatal development during early aerial organ development by the inhibition of *MUTE* and *TMM*, which is consistent with decreased stomatal density under B deficiency (Figure 4e).

In MC, down-regulated genes were enriched in photosynthesis and chloroplast function-related pathways. The down-regulation of these genes due to B deficiency is consistent with the impaired photosynthesis efficiency of MC, which also explains the reduced plant growth rate and biomass accumulation. Photosynthesis-related genes, including *PsPSAN*, *PsPSAL*, *PsPSAF* (photosystem I reaction center), *PsPSB28*, *PsPSBO*, *PsPSBY*, *PsPNSL1* (photosystem II reaction center), were down-regulated. The down-regulation of photosynthesis-related genes is consistent with the mechanism of reduced photosynthesis rate due to B deficiency from a tissue-specific perspective (Figure 4d)(Han et al., 2008).

### Genes involved in meristem and mesophyll cell development

*WUSCHEL-related homeobox 8* (*Psat6g236200*, *PsWOX8*) was identified as a down-regulated DEG in both SAM and MC (Table S4). In addition, *LONELY GUY* (*LOG*) family members *Psat6g021440* (*PsLOG8*) and *Psat6g068040* (*PsLOG2*), which are involved in cytokinin metabolism, were down-regulated in MC and SAM, respectively (Table S4). These observations suggest that SAM activity and MC development might be impaired under B deficiency. After 10 days of B deficiency, the size of *P. sativum* apical buds was significantly reduced, consistent with such a conclusion (Figure 4c). In MC, *P. sativum* homologous genes *Psat6g028400* (*PsKNAT3*) and *Psat0s149g0040* (*PsKNAT4*) from the *KNOX* gene family, which induce leaf formation and development, were down-regulated (Table S4). Auxin-related genes were identified in all cell types and many were induced by B deficiency (Table S4). In these results, we observed that the process of SAM development leading to MC is coherent and highly significant. To further investigate the crucial regulatory factors involved in this process and the role of B, we conducted trajectory construction on specific cell types.

### Cell differentiation trajectory of shoot apical meristem to mesophyll cell

During the development of the SAM, leaf primordia gradually expand and develop into mature leaves. Within this process, a subset of cells enters a proliferative phase, becoming what is referred to as proliferating mesophyll cells (PMCs). These PMCs not only express MC marker genes, but also significantly express genes which have been identified as markers for proliferating cells (Figure 1b, Table S2). Eventually, as the leaf matures, the MCs that cease division transition into mature mesophyll cells (MMCs), at which point they become the primary sites of photosynthesis (Fleming, 2005). To assess the impact of B deficiency on this developmental process, pseudotime analysis was performed on SAM, UM, and MC, with SAM set as the starting point for pseudotime (Figure 5a-b). Both B25 and B0 samples were projected onto the three main branches of the pseudotime analysis, including four trajectory states, with cell differentiation increasing along the two trajectories as pseudotime progressed (Figure 5c, Figure S6d). SAM mainly clustered in state 1, PMC clustered in state 3, and MCs were identified in all the states (Figure S6a-e). Based on the differentiation pattern and the expression of photosynthesis-related genes, cell clusters in state 2 at the late stage of the pseudotime axis are proposed as MMCs (Figure 5c, Figure S6e). Boron deficiency significantly reduced the proportion of PMCs, indicating that B deficiency affects the progress of differentiation of the tissue (Figure S6d-e). By analyzing the gene expression trends over pseudotime, we can decipher the impact of B deficiency on the different fates of SAM cells. Cells were allocated to three main states and two branches, based on the branching point that occurs when differentiating from state 1 to state 2 and state 3 (Figure S6a, c, e). The gene expression changes were categorized according to the trends over pseudotime and on different branches. Based on the similarity of expression trends, five gene clusters were obtained, with cluster 1 and 2 showing branch specificity (Figure 5d, Figure S6f). Cluster 1 genes show an up-regulated expression on the MMC branch as pseudotime progresses, while no significant changes are observed in PMC. Genes in cluster 1 have functions mainly related to DNA packaging, chromatin assembly, nucleosome assembly, and protein-DNA complex assembly pathways (Figure 5d, Table S9). This result indicates that chromatin structural changes mediated by histones play an important role in the gene expression during SAM differentiation and development (Figure 6a-c). Additionally, the 6 histone-related DEGs showing greatest fold difference over pseudotime change are displayed, revealing that these histone genes are highly expressed in the late stage of MC development (Figure 6b).

**Figure 5.**
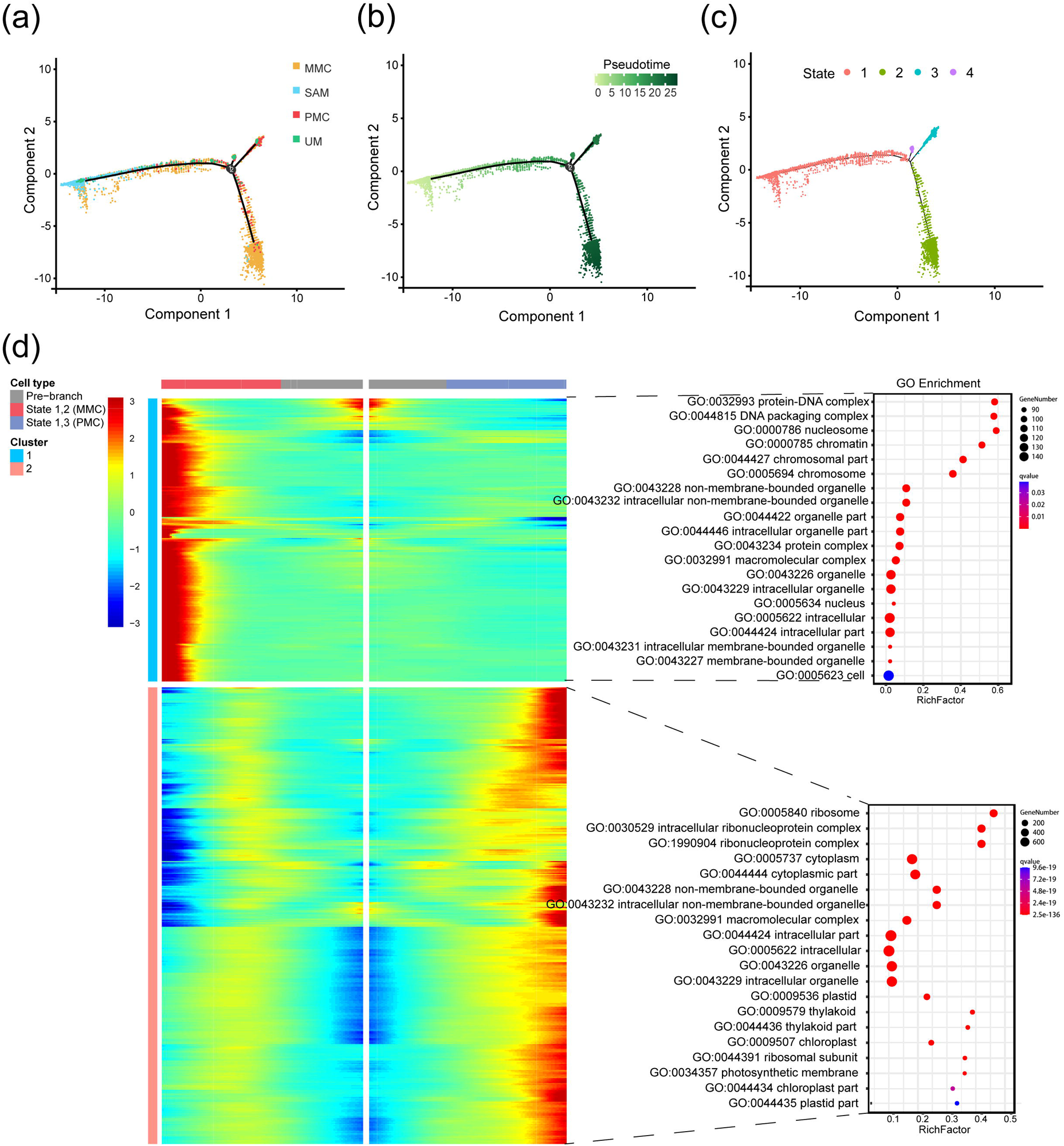
Differentiation trajectory from SAM to MC by pseudotime analysis. (a-c) The distribution of cells along the pseudotime trajectory presented by cell types (a), pseudotime states (b) and branch states (c). (d) Branched heatmap showing DEGs with highly significant branch-specific expression patterns in pseudotime (q values less than 1 × 10^−6^) and significantly enriched GO terms (FDR ≤ 0.05) showing the function of these genes. The detailed information of these genes is given in Table S9. The pre-branch point represents the beginning of pseudotime. Gene expression levels have been normalized by Z-score. Color bar for heatmap indicates the relative expression level. The bubble color (red to violet) in GO enrichment plot indicates the enrichment significance (FDR value) of the GO term, while the bubble size indicates DEG numbers enriched in each GO term.

**Figure 6.**
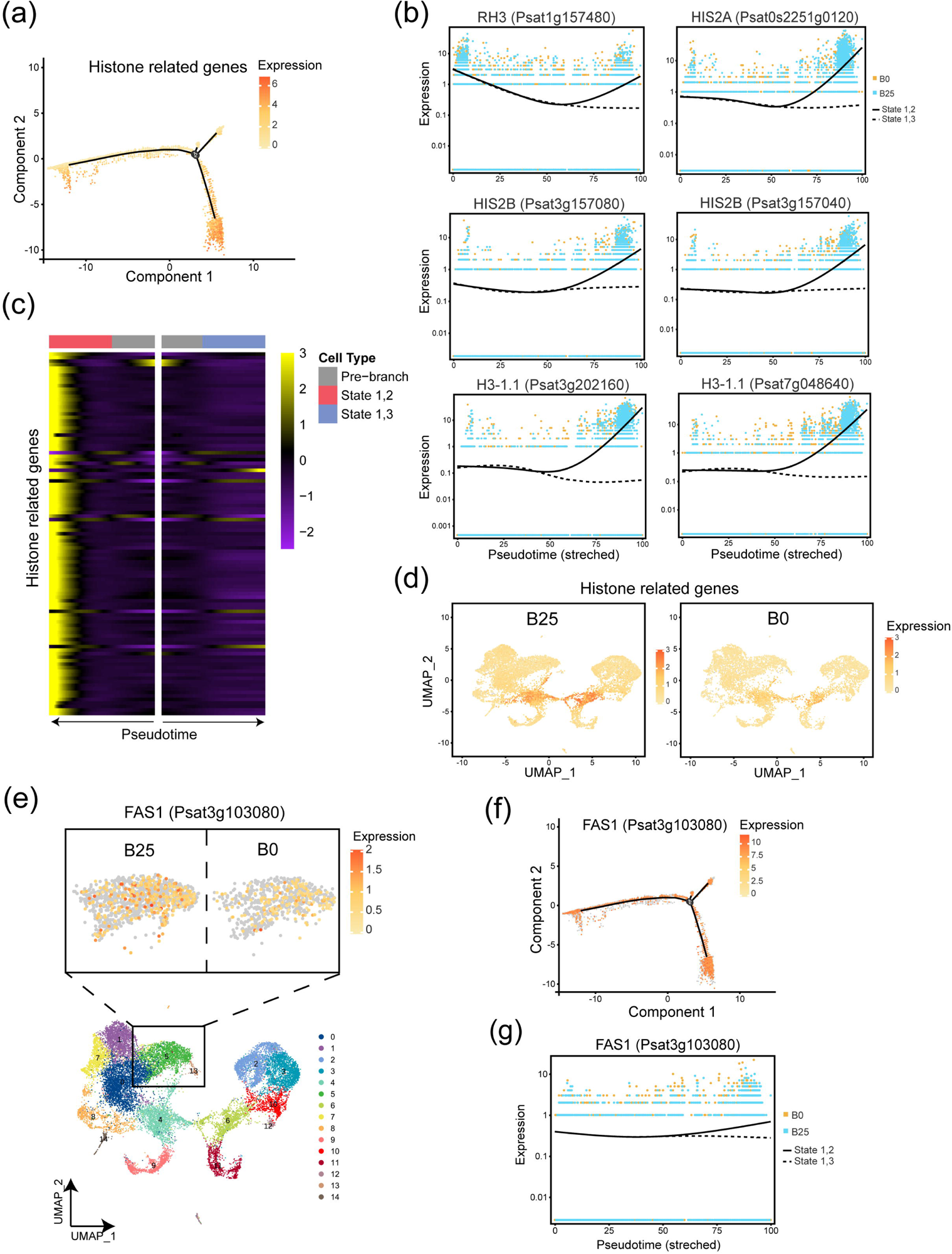
The expression patterns of histone-related genes and chromatin assembly factor subunit *FASCIATA1* in different cell fates and their responses to B deficiency. (a) Expression of histone-related genes along the pseudotime trajectory. Color intensity corresponds to expression level. (b) The expression trends of six strongest histone-related DEGs before and after MC cell differentiation. (c) Heatmap showing all histone-related genes (DEGs) with significant expression trends in different branches. Information on these genes is given in Table S9. (d) UMAP plots showing the down-regulation of the histone related genes under B deficiency. Color intensity corresponds to expression level (e-g) The UMAP and pseudotime expression patterns of *PsFAS1* including down-regulation in SAM (cluster 5) under B deficiency (e), along the pseudotime trajectory (f) and different expression trends on the differentiation trajectory from SAM to MC. Solid lines represent expression from state 1 to state 2. Dashed lines represent expression from state 1 to state 3. Each dot represents an individual cell (blue dots represent cells of B25 while orange dots represent cells of B0).

Boron deficiency was also found to suppress the expression of 83 histone genes (Figure 6d, Table S9), suggesting that B deficiency could induce chromatin structural changes by regulating histone gene expression, with these changes affecting the developmental process and fate of SAM cells. Cluster 2 genes are significantly up-regulated in the PMC branch, with no noticeable differences in MMC. Genes in cluster 2 show functions predominantly related to ribosome structure and function, and peptide biosynthetic processes (Figure 5d, Table S9). Boron deficiency leads to the down-regulation of a large number of such ribosome-related genes (Table S4). Meristem cells have a high protein synthesis activity (Murray et al., 2012). This activity is potentially reduced under B deficiency, potentially impairing cell proliferation.

Thus, we propose that B can regulate molecular pathways determining cell proliferation and cell fate in meristems. This function could explain the necessity of B for meristem function, and in turn provides a molecular explanation for the role of B nutrition in plant growth and development.

### Construction of a transcription factor interaction network

In order to investigate the potential TFs involved in SAM development and the effects of B deficiency on the TF regulatory network, we examined the highly expressed TF genes in different cell types and the effects of B deficiency on their expression. The greatest number of TF genes was identified in SAM (Figure 7a). However, B deficiency induced the expression of the most TF genes in MC. Among which, a large number of *WRKY*-family TF genes were significantly induced by B deficiency, including the previously reported B-responsive *WRKY6* (Table S10)(Kasajima et al., 2010). In addition, we identified the up-regulation of genes encoding TIFY family members, core components of the jasmonic acid signaling pathway, and the NAC family involved in plant growth and development. Some genes encoding members of the ethylene-responsive AP2/ERF family and cytokinin signaling pathway ARR-B family were induced by B deficiency in all cell types, indicating the importance of these TF gene families in response to B deficiency (Table S10). By constructing a protein interaction network of *A. thaliana* homologous TFs affected by B deficiency using STRING (Szklarczyk et al., 2018), we found that *SWI3A* and *SWI3C* interact with most TFs suggesting that they could be core components of a B-responsive TF co-response network (Figure 7b). SWI2A and SWI3C proteins are components of the SWITCH DEFECTIVE/SUCROSE NON-FERMENTABLE (SWI/SNF) chromatin remodeling complex. The SWI/SNF complex regulates gene expression by altering chromatin structure (Bieluszewski et al., 2023). In SAM, these two TF genes not only respond to B deficiency signals but also play a central regulatory role by interacting with other TFs. In addition, *TCP3*, *MYC2*, and *ERF4* also play potentially important regulatory roles (Figure 7b). Our previous bulk RNA-seq study showed that some TFs likely coordinate hormone signaling pathways to influence the response of *P. sativum* shoot apices to B deficiency stress (Chen et al., 2022). The current results further reveal the cell- and tissue-specific responses of TFs to B deficiency (Figure 7c).

**Figure 7.**
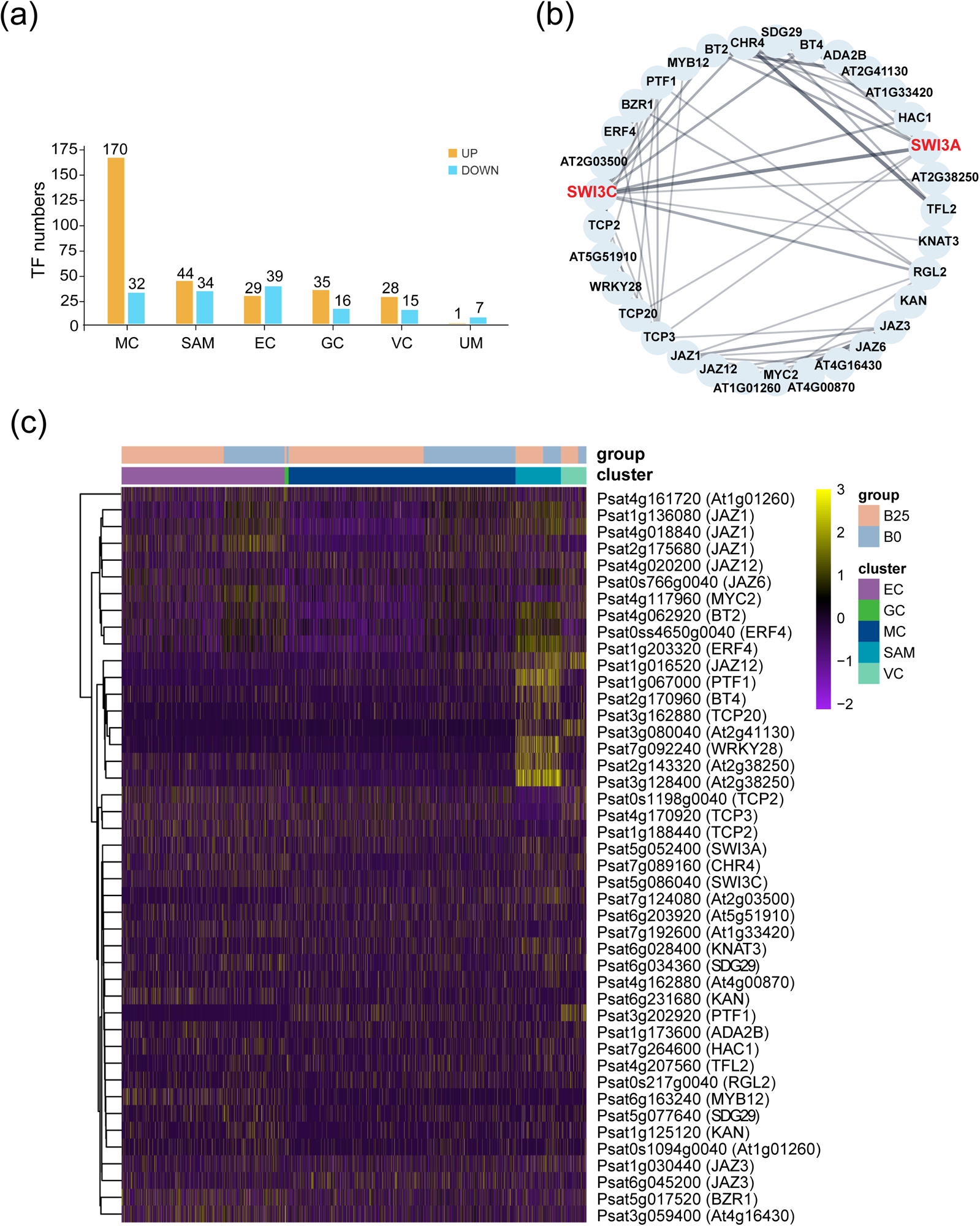
TF network analysis in cell-type-specific responses to B deficiency. (a) Numbers of differentially expressed TF genes in response to B deficiency. Genes upregulated (orange) or downregulated (blue) in B0 relative to B25. Information on these TF genes and their homologs in *A. thaliana* is given in Table S10. (b) Putative protein-protein interaction network of 33 differentially expressed TFs in response to B deficiency. String physical subnetwork was constructed using the homologs in *A. thaliana*. SWI3A and SWI3C, shown in red, are the most-connected proteins in this network. (c) Heatmap showing the cell-type-specific expression of 44 homologous pea genes encoding these core TFs in response to B deficiency (up, yellow; down, purple).

## Discussion

### Application of scRNA-seq to investigate the role of B

Although scRNA-seq has been applied in several studies of plants, its application in nutrient stress is limited. This study revealed cell-type-specific genes in *P. sativum* shoot apices and information on the transcript levels of cell-type-specific responses to B deficiencyIt not only provides additional insights for unraveling cell specificity in the *P. sativum* genome but also identifies potential regulatory factors associated with shoot development in response to B deficiency.

### Identification of cell types and their new markers in *P. sativum* shoot apices

As one of the more widely-studied plants, *P. sativum* shares many similarities with other dicotyledonous plants in its shoot apex. However, compared to *A. thaliana* there is much less information available at the molecular level because genome annotation and functional analysis has been less advanced in P sativum. Furthermore, the lack of cell-type-specific marker genes in legume plants makes cell annotation more challenging. Although scRNA-seq studies of *L. japonicus* root (Sun et al., 2022b) and peanut (*Arachis hypogaea*) leaf (Liu et al., 2021a) have been published previously, no scRNA-seq data has been reported for legume shoot apices. In this study we identify the transcript differences among different cell types in *P. sativum* shoot apices as well as validate cell-type-specific markers using the LCM technology. In addition to validating markers homologous to some previously reported, we also identified new markers for different cell types in *P. sativum*. For example, *AUXIN BINDING PROTEIN 19A* (*PsABP19A*, *Psat1g133880*, *Psat1g133840*) was identified as a new marker for MC. We know that auxin signaling has a regulatory role in the differentiation fate and cell cycle of *A. thaliana* MCs, and the reprogramming of differentiated MCs is regulated by different auxin signals (Sakamoto et al., 2022). This suggests that in *P. sativum*, this gene may be important for MC development-related signaling. *CYTOCHROME P450 96A15* (*PsCYP96A15*, *Psat6g133400*) and *3-KETOACYL-COA SYNTHASE 20* (*PsKCS20*, *Psat6g136720*) were identified as new markers for EC. *AtKCS20* is involved in the synthesis of the important precursor of very-long-chain fatty acids for *A. thaliana* epidermal cuticular wax (Lee et al., 2009). *AtCYP96A15* (also known as *ALKANE HYDROXYLASE MAH1*) is specifically expressed in *A. thaliana* pavement cells, and it also plays an essential role in cuticular wax synthesis in epidermal cells (Greer et al., 2007). In addition to *GSTU8*, which was identified as a SAM marker in *Z. mays* (Satterlee et al., 2020), we also identified another glutathione-related gene, *GLUTAMATE DEHYDROGENASE 3* (*Psat1g096200*, *PsGSH3*), which is significantly up-regulated in SAM. This further suggests that glutathione transferases not only maintain the size and redox balance of root meristems (Horváth et al., 2019) but may also have important regulatory roles in SAM function. Further analysis of gene expression profiles in specific cell types is now possible and will provide valuable knowledge about the characteristics of each cell type, such as in metabolism, differentiation and responses to stress.

### Shoot apex cells exhibit varying responses to B deficiency

Boron has an indispensable role in the structure and function of plant cell walls and membranes. It regulates cell wall formation by cross-linking two rhamnogalacturonan II (RG-II) monomers to form RG-II-B (O’Neill et al., 2001). Moreover, B can affect plasma membrane integrity by forming GIPC-B-RG-II complexes through RG-II and the membrane’s GIPCs (Voxeur and Fry, 2014). One clear phenotype of B deficiency is the inhibition of root elongation (Poza-Viejo et al., 2018). Previous studies have shown that B deficiency affects normal physiological activities in the shoot, including impaired photosynthesis in MC (Figure 4d)(Han et al., 2008), impeding nutrient absorption and transport by affecting the stability of vascular bundles (Pommerrenig et al., 2019), and restricting the development of root meristem (Matthes et al., 2022). However, observing the effects of B deficiency on different cell types in the shoot apex at the single-cell transcript level is a novel aspect of this study.

Boron deficiency does not affect the number of cell clusters predicted informatically, and all identified clusters appear in both B25 and B0 (Figure 4a, f, g). However, most genes respond to B deficiency in a cell-type-specific manner. Among them, the highest number of up-regulated DEGs was identified in MC, while the highest number of down-regulated DEGs was identified in VC (Figure 4b). Previous research has shown that in low-B-demanding rapeseed varieties, mesophyll and vascular growth can be maintained even under B starvation (Pommerrenig et al., 2019). These findings emphasize the plasticity and adaptability of plant mesophyll and vascular cells in coping with B deficiency potentially by regulating gene expression in a cell-type-specific manner. Furthermore, when we applied a higher FC threshold to screen for DEGs, we found that GC, a less abundant cell type, had the most DEGs with FC greater than 2 (Figure S5). This suggests that although MC is more sensitive to moderate gene expression changes induced by B deficiency, GC may have a stronger response by specific genes. Due to the unique function and cell structure of GC, stomata are under rapid and dynamic control in response to abiotic stress (Hedrich and Shabala, 2018). The functions of strongly induced DEGs in GC under B deficiency are focused on cell surface receptor signaling, and ABA-related genes are also identified as significantly up-regulated in GC, indicating that GC responds to B deficiency through ABA signaling, leading to the activation of down-stream signals (Table S4). We also observed a decrease in stomatal density as a result of B deficiency (Figure 4e). These results not only point to the ABA responsive mechanism of stomatal closure caused by B deficiency (Wimmer and Eichert, 2013) but also suggest the possibility of B deficiency affecting stomatal development at an early stage (Wei et al., 2022).

Boron deficiency has a significant impact on the expression of cell wall biosynthesis and degradation-related genes, indicating that plants may maintain cell wall stability and structural integrity by adjusting the expression of cell wall-related genes under B deficiency (Chen et al., 2022). This response also shows some specificity in different cell types. We found that cellulose synthase genes (*CesA*) were up-regulated in most of the cell types under B deficiency, which might reflect a strategy for plants to increase cellulose synthesis to strengthen cell wall structure in response to B deficiency. Polygalacturonase-related genes (*GPs*) were up-regulated in all cell types, suggesting that plants might increase pectin degradation to alleviate the negative effects of B deficiency on cell wall stability. We also observed that cellulose degradation enzyme genes encoding the XTH family (including both xyloglucan endotransglucosylase (XET) and xyloglucan endohydrolase (XEH) enzymes) were up-regulated in EC and GC, which could imply that plants might regulate cellulose degradation to alter cell wall elasticity and stability under B deficiency. Previous bulk RNA-seq results were consistent with B deficiency damaging the structure and function of the cell wall, triggering a series of compensatory mechanisms, leading to plants coping with such stress through biosynthesis and cell wall modification (Chen et al., 2022). The XTH family enzymes, have different effects on the structure of the primary cell wall (Eklöf and Brumer, 2010). As the outermost cells, EC and GC have thicker cell walls and often exhibit rapid responses under stress (Javelle et al., 2011). The regulation of cell wall plasticity in response to B deficiency stress through the XTH family suggests that B deficiency impacts the cell wall integrity signaling in EC and GC (Humphrey et al., 2007).

### The development of mesophyll cell from shoot apical meristem

Since the application of scRNA-seq in plants, researchers have analyzed the single-cell transcriptome profiles of shoot apices in *A. thaliana* (Zhang et al., 2021b), *Z. mays* (Satterlee et al., 2020), and populus (Conde et al., 2022). These studies have demonstrated the high heterogeneity of shoot apical cells in different species, the continuous morphogenesis events in shoot development, and the identification of new regulatory factors in shoot development and physiology. One aim of the present study is to unravel how B deficiency affects apical bud development signals and to analyze the key regulatory factors involved in this process. To this end, we performed pseudotime analysis on SAM, MC, UM, and PMC, creating a differentiation trajectory for *P. sativum* shoot apices (Figure 5a-c). Analyzing and annotating DEGs along different branches helps to identify key genes that determine or reflect cell fate. Moreover, the differences between samples can reveal the role of B in directing cell differentiation and determining the final cell fate. By clustering DEGs affecting the branching of developmental trajectories, we obtained 5 clusters (Figure S6f). We are particularly interested in genes in clusters 1 and 2, showing opposite expression patterns for specific genes in the two branches. Cluster 1 genes are significantly up-regulated in the MMC branch, while their expression levels are maintained in the PMC branch. Functional annotation analysis of the genes in this cluster reveals that their main functions are concentrated in DNA packaging, chromatin assembly, and nucleosome organization. We identified a large number of histone genes, including *H1*, *H2A*, *H2B*, and *H3*, that are significantly up-regulated in the branch determining the fate of MMCs (Figure 6c, Table S9). The organization and regulation of nucleosomes formed by DNA and histone octamers, as well as the regulation of higher-order chromatin condensation, play roles not only in transcription activation and repression but also in differentiation (Li et al., 2007). Chromatin is a dynamic structure that is constantly reshaped to activate or repress specific genes in response to cellular and environmental signals. Chromatin remodeling also plays a crucial role in regulating the *WUS* gene and maintaining the activity of meristematic tissues (Shen and Xu, 2009). By reconstructing cell differentiation trajectories and gene expression trends, we reveal the impact of chromatin structural changes regulated by histone genes on cell fate during *P. sativum* shoot apex differentiation and development.

### Boron is required in meristem development regulated by histone-led chromatin remodeling

In meristem cells, the advantages of epigenetic regulation include the ability to store information, provide stability, yet allow reversibility (Shen and Xu, 2009). As sessile organisms, plants rely heavily on chromatin-based mechanisms and subsequent transcriptional changes for their response mechanisms when faced with inescapable stresses (Probst and Scheid, 2015). For example, iron deficiency increases dissociation of Shk1 binding protein 1 (SKB1/AtPRMT5) from chromatin and reduces histone methylation levels, leading to enhanced transcription of *bHLH* genes and improved iron uptake in *A. thaliana* (Fan et al., 2014). The role of B in maintaining the meristem development has been demonstrated in both shoots and roots (Abreu et al., 2014; Matthes et al., 2022). Based on our previous morphological analysis of shoot apices, B deficiency results in defects in SAM organization (Chen et al., 2022). Here, we propose a chromatin structure-mediated model in which B deficiency leads to changes in chromatin structure and accessibility, promoting the regulation of TFs, which in turn results in reduced meristem activity and impacts cell fate. In *A. thaliana*, CHROMATIN ASSEMBLY FACTOR-1 (CAF-1) consists of three subunits: FASCIATA1 (FAS1), FASCIATA2 (FAS2), and MULTICOPY SUPPRESSOR OF IRA1 (MSI1) (Shen and Xu, 2009). FAS1 participates in limiting *WUS* expression in the OC region, and *PsFAS1* plays a regulatory role in determining cell differentiation fate, which is inhibited in SAM by B deficiency (Figure 6e-g). By elucidating the role of *PsFAS1* on regulating chromatin structure and identifying its response to B deficiency in SAM, our findings shed light on the mechanistic understanding of how B deficiency influences chromatin remodeling during shoot development. This chromatin structure-mediated model provides insights into the molecular processes underlying the effects of B deficiency on SAM organization and cell fate determination. Further research is warranted to unravel the precise molecular mechanisms and signaling pathways involved in this intricate regulatory network.

### Potential role of SWI/SNF as a core transcription factor in response to B deficiency

TF regulatory networks also play a crucial role in shoot apex development. Key TFs identified in bulk RNA-seq results, such as *PsWRKY40* and *PsERF4*, were also captured by scRNA-seq results (Figure 7b, Table S10). However, they were induced by B deficiency in different cell types. *PsWRKY40* was induced in MC, while *PsERF4* was induced in GC and VC, indicating that the expression of B deficiency-induced TFs is cell-type-specific. Members of the SWI/SNF complex, *SWI3A* and *SWI3C*, significantly responded to B deficiency in SAM and potentially act as core genes in the TF network, coordinating other TFs to regulate downstream signals. The SWI/SNF chromatin remodeling complex alters the contact between DNA and nucleosomes by utilizing ATP to make chromatin structures looser or tighter, thus affecting the accessibility of TFs and other regulatory proteins to specific genes (Bieluszewski et al., 2023). *WUS* expression in the OC region is activated by the SWI/SNF-type ATPase SYD (Shen and Xu, 2009). Our results suggest that SWI/SNF participates in the gene regulatory network under B deficiency.

## Conclusion

This study demonstrates the power of the single-cell transcriptomics approach and reveals strategies by which plants adapt to conditions of B deficiency. We show that responses of *P. sativum* shoot apices to B deficiency show highly pronounced heterogeneity, with distinct cell types exhibiting varying responses that correlated with their specific functions. The pronounced response observed in the GCs highlights their vital role in both withstanding external pressures and participating in signal transduction. Secondly, through pseudotime analysis, we have constructed a developmental trajectory from SAM to MC and shown that chromatin structural changes, regulated by histones and FAS1, may serve as a crucial regulatory pathway governing the progression of SAM development. By providing a precise transcriptome-level analysis, this study offers valuable insights for enhancing plant stress tolerance and for developing genotypes capable of coping with B stress not only in pea but also in a broad range of legume species.

## Materials and methods

### Plant growth and treatment

Pea (*P. sativum*. cv. Caméor) seeds were treated with a 7.5% (w/v) sodium hypochlorite solution for 30 min, followed by soaking in a 0.5 mM CaCl_2_ at 25 °C in the dark for 8 h. Germination took place over 48 h in an aeroponic system containing 0.5 mM CaCl_2_. Seedlings with roots 3-4 cm in length were then cultivated at 25 °C in a one-fourth-strength modified Hoagland solution (pH 5.5), under a 16 h light (100 µmol photons m^−2^s^−1^) and 8 h dark cycle at 75% relative humidity. The B-free nutrient solution (B0) was prepared by removing B from water using an ion exchange resin (Amberlite IRA743 free base, Sigma-Aldrich, St. Louis, MO, USA) over 3 d. The B25 nutrient solution was prepared by adding 25 μM H_3_BO_3_ to the base solution. After 10 d of hydroponic growth, the shoot apices were sampled for immediate protoplast isolation.

### Protoplast preparation

Fifty shoot apices were cut into 1-2 mm strips and placed in a filtered enzyme solution containing 10 ml of 1.5% (w/v) cellulose R10, 1.5% (w/v) macerozyme R10, 0.5% (w/v) pectinase, 8% (w/v) mannitol (without Ca^2+^ and Mg^2+^), 0.02 M KCl, 0.01 M MES (pH 5.7), 0.01 M CaCl_2_, and 0.25% (w/v) BSA. The centrifuge tube containing the enzyme solution and tissue was placed on an orbital shaker at 30°C with a shaking speed of 75 rpm in the dark for 1 h. The tissue was then washed twice with 8% (w/v) mannitol and filtered through a 40 μm cell strainer twice. A 5 µl protoplast suspension was mixed with 5 µl of 0.4% (w/v) trypan blue dye, and cell concentration and viability were measured using a hemocytometer and a light microscope. Finally, the protoplast suspension concentration was adjusted to 1000-2000 cells/µl using an 8% (w/v) mannitol solution in preparation for loading onto the chromium controller of the 10x Genomics platform (10x Genomics, Pleasanton, CA, USA).

### scRNA-seq library construction

Approximately 2×10^4^ isolated single cells and enzyme gel-beads were packed into a single oil droplet for scRNA-seq library construction. The scRNA-seq libraries were constructed using the Chromium Single-Cell 3’ GEM (Gel Beads-in-Emulsion) Library & Gel Bead Kit v3, following the user manual’s instructions (10x Genomics, Pleasanton, CA, USA). The Single-Cell 3’ Library protocol generated standard Illumina paired-end constructs. The libraries were quality checked using the High Sensitivity DNA assay Kit (Agilent Technologies, Santa Clara, CA, USA). Finally, RNA quantification was performed using the ABI StepOnePlus Real Time PCR System (Thermo Fisher Scientific, Waltham, MA, USA), followed by high-throughput sequencing on the Illumina HiSeq2500. The 10X Genomics Cell Ranger software (10x Genomics, v3.1.0) was used for alignment and quantification. Cell Ranger employed the STAR (Spliced Transcripts Alignment to a Reference) aligner to map single-cell sequencing reads to the reference genome, *P. sativum* v1a (Kreplak et al., 2019). After filtering and correcting barcodes and UMIs, unique UMI counts and cell barcodes were used to generate gene-by-cell matrices for downstream analysis.

### Data processing and cell clustering

First, the DoubletFinder was used to remove GEMs with doublets (McGinnis et al., 2019). The real doublets were distinguished from singlets by identifying real cells with high proportions of artificial neighbors in gene expression space. The low-quality cells were filtered out by setting cutoff values for the number of expressed genes per cell to greater than 500 and less than 8000. To visualize the data, dimension reduction and clustering analysis were performed on the raw count matrices using the Seurat (v3.1.1) package in R (v4.0.2). The data were reduced using PCA, and based on the classification results of cell subpopulations, UMAP was further employed to visualize the cell subpopulation classifications, resulting in 15 cell clusters. To identify these cell types, marker genes reported in *A. thaliana* and *Z. mays* were used, and *P. sativum* marker genes obtained through orthologous gene comparison (Table S2). *A. thaliana* protein sequences were downloaded from TAIR (Garcia-Hernandez et al., 2002). In this way six cell types were identified. Additionally, marker genes for each cell cluster were identified using the Wilcoxon Rank Sum test (with default parameters: likelihood ratio test) through the FindAllMarkers in Seurat. The filtering criteria for cell cluster-specific up-regulated genes were: genes expressed in more than 25% of cells in the target or control cluster, *P*-value ≤ 0.01; and gene expression log_2_FC ≥ 0.36. When identifying DEGs within different cell types induced by B deficiency, we used a threshold of |log2FC| ≥ 0.36 and *P* < 0.05.

### Gene enrichment analysis

All DEGs were assigned to GO terms in the GO database (http://www.geneontology.org/) and to pathway terms in the KEGG database (http://www.genome.jp/kegg/). A hypergeometric test was used to identify enriched GO and KEGG terms. GO and KEGG terms that were overrepresented and had a Benjamini-Hochberg adjusted *p*-value of less than 0.05 were included in subsequent analyses.

### Laser capture microdissection and qRT-PCR

LCM was performed following existing methods (Takahashi et al., 2010). Pea shoot apices were embedded in paraffin (Paraplast Xtra; Fisher Scientific, Pittsburgh, PA, USA) using a previously reported microwave-based method (Takahashi et al., 2010). After slicing (10 μm thick) and transferring to microscope slides (PEN membrane glass slide, Thermo Fisher Scientific, Waltham, MA, USA) and drying, MC, SAM, and vascular bundle cells were collected from different tissue sections using a Zeiss PALM MicroBeam Laser Microdissection System (Carl Zeiss, Jena, Germany). Each cell type had four replicates, and more than 300 sections were captured for each replicate. Subsequently, total RNA extraction was performed using a Pico-Pure™ RNA isolation kit (Thermo Fisher Scientific, Sunnyvale, CA, USA). The extracted total RNA was quantified using Quant-iT™ RiboGreen RNA reagent and kit (Invitrogen, Carlsbad, CA, USA). The quality of the total RNA extracted from target cells was evaluated using the RNA 6000 Pico kit on an Agilent 2100 Bioanalyzer (Agilent Technologies, Santa Clara, CA, USA). RNA integrity was determined based on the RNA Integrity Number (RIN) (Schroeder et al., 2006) and analyzed using 2100 Expert software (version B.02.02, Eukaryote Total RNA Pico Mode, Agilent Technologies, Santa Clara, CA, USA). The RIN of each tissue was above 6, which meets the criteria for qRT-PCR analysis. The expression of marker genes was detected by qRT-PCR using the One-step TB Green PrimeScriptTM RT-PCR kit II (Takara, Osaka, Japan). All genes were normalized against the level of an *actin* reference gene (Foo et al., 2005). Primer sequences are given in Table S11.

### Pseudotime trajectory analysis

Pseudotime analysis was performed using Monocle (v3.0) to construct cell matrices and gene expression profiles, visualizing developmental trajectories while preserving the fundamental relationships among cell types in reduced dimensions (Trapnell et al., 2014). Monocle can utilize gene expression level signals in all cells and based on the pseudotime values of each cell, screen DEGs along the timeline to identify key genes related to developmental differentiation processes. The developmental trajectory was determined from SAM to MC and two distinct branches identified. By clustering genes with similar expression trends, we identified DEGs that changed along the developmental trajectory. Their potential biological functions were further analyzed through GO and KEGG enrichment.

### Transcription factor interaction network construction

To analyze core TFs, all expressed TFs in the samples were annotated using the plant TF database PlantTFDB (http://planttfdb.cbi.pku.edu.cn) (Guo et al., 2007). Differentially expressed TFs (|log2FC| ≥ 0.36 and *P* ≤ 0.01) were screened and B-responsive highly expressed TFs connected in the protein-protein interaction network using the STRING database. Finally, Pearson correlation coefficients (PCCs) were calculated between differentially expressed TFs and a co-expression network constructed using Cytoscape_v3.7.2 (PCCs ≥ 0.6, *P* < 0.05).

### Data analysis and visualization

Data were analyzed using SPSS Statistics 20.0 (SPSS Inc., Chicago, IL). Duncan’s least significant difference test was performed through analysis of variance (ANOVA) to determine the significance at *P* < 0.05. Visualization tools mainly used included Adobe Illustrator, R software, and OmicShare tools (https://www.omicshare.com/tools) (Gene Denovo Biotech. Co., Guangzhou, China).

## Date availability

Single-cell RNA-seq data have been deposited in the Sequence Read Archive (SRA) database at the National Center for Biotechnology Information (NCBI) under the accession number PRJNA983513.

## Author contributions

X.C., S.M.S. and M.Y. conceptualized and supervised the research. X.C. and Y.R. prepared the materials for scRNA-seq. X.C. performed data analysis. X.C., H.T. and M.N preformed and revised the LCM part. X.C. wrote the manuscript. X.C., S.S., S.M.S. and M.Y. revised the manuscript.

## Acknowledgements

This work was supported by the National Natural Science Foundation of China (Grant No. 32172672,31672228, 31172038), Ministry of Science and Technology of China (CB02-07, WQ20174400441, 2018YFD0201203, G2021030008L, G2021030014L, DL20200230002), the Science and Technology Department of Guangdong Province (Grant No. 2018A050506085, 2015A040404048, 163-2018-XMZC-0001-05-0049, 2022B1212010015 2017-1649), the Higher Education Department of Guangdong Province (Grant No. 2020KCXTD025), MEXT KAKENHI (Grant No. JP20H05912) and the Australian Research Council (Grant No. CE200100015).

## Conflict of interest

The authors declare that there is no conflict of interest.

## Legends of supporting information

**Figure S1** Overview of scRNA-seq analysis. (a) Protoplast preparations for scRNA-seq. (b) Venn diagram showing numbers of unique and common genes detected in bulk RNA-seq and scRNA-seq in B0 and B25 samples.

**Figure S2** Numbers of cells in each scRNA-seq cluster and correlation of each cluster. (a) Number of cells in each cluster in B0 and B25 samples. (b) Correlations between 15 cell clusters expressed as proportion of detected genes in common (greater than 0.5, red; less than 0.5, blue).

**Figure S3** KEGG enrichment annotation of genes expressed specifically in the five known cell types. (a) SAM (b) MC (c) EC (d) GC (e) VC.

**Figure S4** GO term enrichment annotation of genes expressed specifically in the five known cell types. (a) SAM (b)MC (c) EC (d) GC (e) VC.

**Figure S5** Numbers of DEGs in response to B-deficiency in six cell types. DEGs were identified as up (orange) or down (blue) in B0 relative to B25 with a fold-change of |log_2_FC| ≥ 1.

**Figure S6** Pseudotime trajectory from SAM to MC. (a-c) The distribution of cells along the pseudotime trajectory color-coded by cell type (a), pseudotime states, represented by intensity of green (b), and branch states, color-coded (c). MMC, mature mesophyll cells; SAM, shoot apical meristem cells; PMC, proliferating mesophyll cells; UM, unknown meristem. (d-e) Cell proportion of four states in B0 and B25 samples (d), and four different cell types (e). (f) Heatmap showing all gene clusters over the pseudotime trajectories.

**Table S1.** Single cell sequencing results.

**Table S2.** Marker genes used for cluster annotation.

**Table S3**. Cell numbers change after boron deficiency (BD).

**Table S4.** Differentially expressed genes (DEGs) expression changes in different cell types.

**Table S5**. KEGG significant enrichment for DEGs from Table S4.

**Table S6.** GO significant terms for DEGs from Table S4.

**Table S7**. Differentially expressed boron transporters in different cell types.

**Table S8.** Cell wall related DEGs expression in different cell types.

**Table S9**. Differentiation fate DEGs from pseudotime analysis (cluster1 & cluster2).

**Table S10.** Transcription factors (TFs) expression changes in different cell types.

**Table S11**. Primer sequences used for qRT-PCR in laser capture microdissection (LCM).

## Notes

### Competing Interest Statement

The authors have declared no competing interest.

## References

Abreu, I., Poza, L., Bonilla, I. and Bolaños, L. (2014) Boron deficiency results in early repression of a cytokinin receptor gene and abnormal cell differentiation in the apical root meristem of *Arabidopsis thaliana*. Plant Physiol. Biochem. 77, 117–121.

Bai, Y., Liu, H., Lyu, H., Su, L., Xiong, J. and Cheng, Z.M. (2022) Development of a single-cell atlas for woodland strawberry (Fragaria vesca) leaves during early Botrytis cinerea infection using single cell RNA-seq. Hort. Res. 9, uhab055.

Berkowitz, O., Xu, Y., Liew, L.C., Wang, Y., Zhu, Y., Hurgobin, B., Lewsey, M.G. and Whelan, J. (2021) RNA-seq analysis of laser microdissected *Arabidopsis thaliana* leaf epidermis, mesophyll and vasculature defines tissue-specific transcriptional responses to multiple stress treatments. Plant J. 107, 938–955.

Bezrutczyk, M., Zöllner, N.R., Kruse, C.P.S., Hartwig, T., Lautwein, T., Köhrer, K., Frommer, W.B. and Kim, J.Y. (2021) Evidence for phloem loading via the abaxial bundle sheath cells in maize leaves. Plant Cell 33, 531–547.

Bieluszewski, T., Prakash, S., Roulé, T. and Wagner, D. (2023) The role and activity of SWI/SNF chromatin remodelers. Annu. Rev. Plant Biol. 74, 28.21–28.25.

Bowman, J.L. and Eshed, Y. (2000) Formation and maintenance of the shoot apical meristem. Trends Plant Sci. 5, 110–115.

Carles, C.C. and Fletcher, J.C. (2003) Shoot apical meristem maintenance: the art of a dynamic balance. Trends Plant Sci. 8, 394–401.

Chen, X., Humphreys, J.L., Ru, Y., He, Y., Wu, F., Mai, J., Li, M., Li, Y., Shabala, S., Yu, M. and Smith, S.M. (2022) Jasmonate signaling and remodeling of cell wall metabolism induced by boron deficiency in pea shoots. Environ. Exp. Bot. 201, 104947.

Chen, Y., Tong, S., Jiang, Y., Ai, F., Feng, Y., Zhang, J., Gong, J., Qin, J., Zhang, Y., Zhu, Y., Liu, J. and Ma, T. (2021) Transcriptional landscape of highly lignified poplar stems at single-cell resolution. Genome Biol. 22, 319.

Conde, D., Triozzi, P.M., Pereira, W.J., Schmidt, H.W., Balmant, K.M., Knaack, S.A., Redondo-López, A., Roy, S., Dervinis, C. and Kirst, M. (2022) Single-nuclei transcriptome analysis of the shoot apex vascular system differentiation in *Populus*. Development 149, dev200632.

Denyer, T., Ma, X., Klesen, S., Scacchi, E., Nieselt, K. and Timmermans, M.C.P. (2019) Spatiotemporal developmental trajectories in the *Arabidopsis* root revealed using high-throughput single-cell RNA sequencing. Dev. Cell 48, 840–852.

Eklöf, J.M. and Brumer, H. (2010) The *XTH* gene family: an update on enzyme structure, function, and phylogeny in xyloglucan remodeling. Plant Physiol. 153, 456–466.

Fan, H., Zhang, Z., Wang, N., Cui, Y., Sun, H., Liu, Y., Wu, H., Zheng, S., Bao, S. and Ling, H.Q. (2014) SKB1/PRMT5-mediated histone H4R3 dimethylation of Ib subgroup *bHLH* genes negatively regulates iron homeostasis in *Arabidopsis thaliana*. Plant J. 77, 209–221.

Fleischer, A., O’Neill, M.A. and Ehwald, R. (1999) The pore size of non-graminaceous plant cell walls is rapidly decreased by borate ester cross-linking of the pectic polysaccharide rhamnogalacturonan II. Plant Physiol. 121, 829–838.

Fleming, A.J. (2005) The co-ordination of cell division, differentiation and morphogenesis in the shoot apical meristem: a perspective. J. Exp. Bot. 57, 25–32.

Foo, E., Bullier, E., Goussot, M., Foucher, F., Rameau, C. and Beveridge, C.A. (2005) The branching gene *RAMOSUS1* mediates interactions among two novel signals and auxin in pea. Plant Cell 17, 464–474.

Gala, H.P., Lanctot, A., Jean-Baptiste, K., Guiziou, S., Chu, J.C., Zemke, J.E., George, W., Queitsch, C., Cuperus, J.T. and Nemhauser, J.L. (2021) A single-cell view of the transcriptome during lateral root initiation in *Arabidopsis thaliana*. Plant Cell 33, 2197–2220.

Garcia-Hernandez, M., Berardini, T., Chen, G., Crist, D., Doyle, A., Huala, E., Knee, E., Lambrecht, M., Miller, N., Mueller, L.A., Mundodi, S., Reiser, L., Rhee, S.Y., Scholl, R., Tacklind, J., Weems, D.C., Wu, Y., Xu, I., Yoo, D., Yoon, J. and Zhang, P. (2002) TAIR: a resource for integrated *Arabidopsis* data. Funct. Integr. Genomics 2, 239–253.

Goldbach, H.E. and Wimmer, M.A. (2007) Boron in plants and animals: Is there a role beyond cell-wall structure? J. Plant Nutr. Soil Sci. 170, 39–48.

Greer, S., Wen, M., Bird, D., Wu, X., Samuels, L., Kunst, L. and Jetter, R. (2007) The cytochrome P450 enzyme CYP96A15 is the midchain alkane hydroxylase responsible for formation of secondary alcohols and ketones in stem cuticular wax of *Arabidopsis*. Plant Physiol. 145, 653–667.

Guo, A.Y., Chen, X., Gao, G., Zhang, H., Zhu, Q.H., Liu, X.C., Zhong, Y.F., Gu, X., He, K. and Luo, J. (2007) PlantTFDB: a comprehensive plant transcription factor database. Nucleic Acids Res. 36, D966–D969.

Han, S., Chen, L.S., Jiang, H.X., Smith, B.R., Yang, L.T. and Xie, C.Y. (2008) Boron deficiency decreases growth and photosynthesis, and increases starch and hexoses in leaves of citrus seedlings. J. Plant Physiol. 165, 1331–1341.

Hedrich, R. and Shabala, S. (2018) Stomata in a saline world. Curr. Opin. Plant Biol. 46, 87–95.

Horváth, E., Bela, K., Holinka, B., Riyazuddin, R., Gallé, Á., Hajnal, Á., Hurton, Á., Fehér, A. and Csiszár, J. (2019) The *Arabidopsis* glutathione transferases, AtGSTF8 and AtGSTU19 are involved in the maintenance of root redox homeostasis affecting meristem size and salt stress sensitivity. Plant Sci. 283, 366–374.

Humphrey, T.V., Bonetta, D.T. and Goring, D.R. (2007) Sentinels at the wall: cell wall receptors and sensors. New Phytol. 176, 7–21.

Javelle, M., Vernoud, V., Rogowsky, P.M. and Ingram, G.C. (2011) Epidermis: the formation and functions of a fundamental plant tissue. New Phytol. 189, 17–39.

Kasajima, I., Ide, Y., Yokota Hirai, M. and Fujiwara, T. (2010) WRKY6 is involved in the response to boron deficiency in *Arabidopsis thaliana*. Physiol. Plant. 139, 80–92.

Kreplak, J., Madoui, M.-A., Cápal, P., Novák, P., Labadie, K., Aubert, G., Bayer, P.E., Gali, K.K., Syme, R.A., Main, D., Klein, A., Bérard, A., Vrbová, I., Fournier, C., d’Agata, L., Belser, C., Berrabah, W., Toegelová, H., Milec, Z., Vrána, J., Lee, H., Kougbeadjo, A., Térézol, M., Huneau, C., Turo, C.J., Mohellibi, N., Neumann, P., Falque, M., Gallardo, K., McGee, R., Tar’an, B., Bendahmane, A., Aury, J.-M., Batley, J., Le Paslier, M.-C., Ellis, N., Warkentin, T.D., Coyne, C.J., Salse, J., Edwards, D., Lichtenzveig, J., Macas, J., Doležel, J., Wincker, P. and Burstin, J. (2019) A reference genome for pea provides insight into legume genome evolution. Nat. Genet. 51, 1411–1422.

Lee, S.B., Jung, S.J., Go, Y.S., Kim, H.U., Kim, J.K., Cho, H.J., Park, O.K. and Suh, M.C. (2009) Two *Arabidopsis* 3-ketoacyl CoA synthase genes, *KCS20* and KCS2/DAISY, are functionally redundant in cuticular wax and root suberin biosynthesis, but differentially controlled by osmotic stress. Plant J. 60, 462–475.

Li, B., Carey, M. and Workman, J.L. (2007) The role of chromatin during transcription. Cell 128, 707–719.

Li, X., Zhang, X., Gao, S., Cui, F., Chen, W., Fan, L. and Qi, Y. (2022) Single-cell RNA sequencing reveals the landscape of maize root tips and assists in identification of cell type-specific nitrate-response genes. Crop J. 10, 1589–1600.

Liang, X., Ma, Z., Ke, Y., Wang, J., Wang, L., Qin, B., Tang, C., Liu, M., Xian, X., Yang, Y., Wang, M. and Zhang, Y. (2023) Single-cell transcriptomic analyses reveal cellular and molecular patterns of rubber tree response to early powdery mildew infection. Plant Cell Environ., 1–16.

Liu, H., Hu, D., Du, P., Wang, L., Liang, X., Li, H., Lu, Q., Li, S., Liu, H., Chen, X., Varshney, R.K. and Hong, Y. (2021a) Single-cell RNA-seq describes the transcriptome landscape and identifies critical transcription factors in the leaf blade of the allotetraploid peanut (*Arachis hypogaea* L.). Plant Biotechnol. J. 19, 2261–2276.

Liu, Q., Liang, Z., Feng, D., Jiang, S., Wang, Y., Du, Z., Li, R., Hu, G., Zhang, P., Ma, Y., Lohmann, J.U. and Gu, X. (2021b) Transcriptional landscape of rice roots at the single-cell resolution. Mol. Plant 14, 384–394.

Liu, Z., Kong, X., Long, Y., Liu, S., Zhang, H., Jia, J., Cui, W., Zhang, Z., Song, X., Qiu, L., Zhai, J. and Yan, Z. (2023) Integrated single-nucleus and spatial transcriptomics captures transitional states in soybean nodule maturation. Nat. Plants 9, 515–524.

Matthes, M.S., Darnell, Z., Best, N.B., Guthrie, K., Robil, J.M., Amstutz, J., Durbak, A. and McSteen, P. (2022) Defects in meristem maintenance, cell division, and cytokinin signaling are early responses in the boron deficient maize mutant *tassel-less1*. Physiol. Plant. 174, e13670.

McGinnis, C.S., Murrow, L.M. and Gartner, Z.J. (2019) DoubletFinder: doublet detection in single-cell RNA sequencing data using artificial nearest neighbors. Cell Syst. 8, 329–337.

Murray, J.A.H., Jones, A., Godin, C. and Traas, J. (2012) Systems analysis of shoot apical meristem growth and development: integrating hormonal and mechanical signaling. Plant Cell 24, 3907–3919.

O’Neill, M.A., Eberhard, S., Albersheim, P. and Darvill, A.G. (2001) Requirement of borate cross-linking of cell wall rhamnogalacturonan II for *Arabidopsis* growth. Science 294, 846–849.

O’Neill, M.A., Warrenfeltz, D., Kates, K., Pellerin, P., Doco, T., Darvill, A.G. and Albersheim, P. (1996) Rhamnogalacturonan-II, a pectic polysaccharide in the walls of growing plant cell, forms a dimer that is covalently cross-linked by a borate ester. J. Biol. Chem. 271, 22923–22930.

Pommerrenig, B., Eggert, K. and Bienert, G.P. (2019) Boron deficiency effects on sugar, ionome, and phytohormone profiles of vascular and non-vascular leaf tissues of common plantain (*Plantago major* L.). Int. J. Mol. Sci. 20, 3882.

Poza-Viejo, L., Abreu, I., González-García, M.P., Allauca, P., Bonilla, I., Bolaños, L. and Reguera, M. (2018) Boron deficiency inhibits root growth by controlling meristem activity under cytokinin regulation. Plant Sci. 270, 176–189.

Probst, A.V. and Scheid, O.M. (2015) Stress-induced structural changes in plant chromatin. Curr. Opin. Plant Biol. 27, 8–16.

Sakamoto, Y., Kawamura, A., Suzuki, T., Segami, S., Maeshima, M., Polyn, S., De Veylder, L. and Sugimoto, K. (2022) Transcriptional activation of auxin biosynthesis drives developmental reprogramming of differentiated cells. Plant Cell 34, 4348–4365.

Satterlee, J.W., Strable, J. and Scanlon, M.J. (2020) Plant stem-cell organization and differentiation at single-cell resolution. Proc. Natl. Acad. Sci. USA 117, 33689–33699.

Schroeder, A., Mueller, O., Stocker, S., Salowsky, R., Leiber, M., Gassmann, M., Lightfoot, S., Menzel, W., Granzow, M. and Ragg, T. (2006) The RIN: an RNA integrity number for assigning integrity values to RNA measurements. BMC Mol. Biol. 7, 3.

Shaw, R., Tian, X. and Xu, J. (2020) Single-cell transcriptome analysis in plants: advances and challenges. Mol. Plant 14, 115–126.

Shen, W.H. and Xu, L. (2009) Chromatin remodeling in stem cell maintenance in *Arabidopsis thaliana*. Mol. Plant 2, 600–609.

Smýkal, P., Aubert, G., Burstin, J., Coyne, C.J., Ellis, N.T.H., Flavell, A.J., Ford, R., Hýbl, M., Macas, J., Neumann, P., McPhee, K.E., Redden, R.J., Rubiales, D., Weller, J.L. and Warkentin, T.D. (2012) Pea (*Pisum sativum* L.) in the genomic era. Agronomy 2, 74–115.

Sun, X., Feng, D., Liu, M., Qin, R., Li, Y., Lu, Y., Zhang, X., Wang, Y., Shen, S., Ma, W. and Zhao, J. (2022a) Single-cell transcriptome reveals dominant subgenome expression and transcriptional response to heat stress in Chinese cabbage. Genome Biol. 23, 262.

Sun, Z., Jiang, S., Wang, D., Li, L., Liu, B., Ran, Q., Hu, L., Xiong, J., Tang, Y., Gu, X., Wu, Y. and Liang, Z. (2022b) Single-cell RNA-seq of *Lotus japonicus* provide insights into identification and function of root cell types of legume. J. Integr. Plant Biol. 65, 1147–1152.

Szklarczyk, D., Gable, A.L., Lyon, D., Junge, A., Wyder, S., Huerta-Cepas, J., Simonovic, M., Doncheva, N.T., Morris, J.H., Bork, P., Jensen, L.J. and Mering, Christian v. (2018) STRING v11: protein–protein association networks with increased coverage, supporting functional discovery in genome-wide experimental datasets. Nucleic Acids Res. 47, D607–D613.

Takahashi, H., Kamakura, H., Sato, Y., Shiono, K., Abiko, T., Tsutsumi, N., Nagamura, Y., Nishizawa, N.K. and Nakazono, M. (2010) A method for obtaining high quality RNA from paraffin sections of plant tissues by laser microdissection. J. Plant Res. 123, 807–813.

Tassoni, A., Tedeschi, T., Zurlini, C., Cigognini, I.M., Petrusan, J.-I., Rodríguez, Ó., Neri, S., Celli, A., Sisti, L., Cinelli, P., Signori, F., Tsatsos, G., Bondi, M., Verstringe, S., Bruggerman, G. and Corvini, P.F.X. (2020) State-of-the-art production chains for peas, beans and chickpeas - valorization of agro-industrial residues and applications of derived extracts. Molecules 25, 1383.

Trapnell, C., Cacchiarelli, D., Grimsby, J., Pokharel, P., Li, S., Morse, M., Lennon, N.J., Livak, K.J., Mikkelsen, T.S. and Rinn, J.L. (2014) The dynamics and regulators of cell fate decisions are revealed by pseudotemporal ordering of single cells. Nat. Biotechnol. 32, 381–386.

Voxeur, A. and Fry, S.C. (2014) Glycosylinositol phosphorylceramides from *Rosa* cell cultures are boron-bridged in the plasma membrane and form complexes with rhamnogalacturonan II. Plant J. 79, 139–149.

Wang, Q., Wu, Y., Peng, A., Cui, J., Zhao, M., Pan, Y., Zhang, M., Tian, K., Schwab, W. and Song, C. (2022) Single-cell transcriptome atlas reveals developmental trajectories and a novel metabolic pathway of catechin esters in tea leaves. Plant Biotechnol. J. 20, 2089–2106.

Wang, Y., Huan, Q., Li, K. and Qian, W. (2021) Single-cell transcriptome atlas of the leaf and root of rice seedlings. J. Genet. Genomics 48, 881–898.

Warington, K. (1923) The effect of boron acid and borax on the broad bean and certain other plants. Ann. Bot. 37, 457–466.

Wei, R., Huang, M., Huang, D., Zhou, J., Pan, X. and Zhang, W.E. (2022) Growth, gas exchange, and boron distribution characteristics in two grape species plants under boron deficiency condition. Horticulturae 8, 374.

Wimmer, M.A. and Eichert, T. (2013) Review: Mechanisms for boron deficiency-mediated changes in plant water relations. Plant Sci. 203–204, 25-32.

Wong, C.E., Bhalla, P.L., Ottenhof, H. and Singh, M.B. (2008) Transcriptional profiling of the pea shoot apical meristem reveals processes underlying its function and maintenance. BMC Plant Biol. 8, 73.

Yang, T., Liu, R., Luo, Y., Hu, S., Wang, D., Wang, C., Pandey, M.K., Ge, S., Xu, Q., Li, N., Li, G., Huang, Y., Saxena, R.K., Ji, Y., Li, M., Yan, X., He, Y., Liu, Y., Wang, X., Xiang, C., Varshney, R.K., Ding, H., Gao, S. and Zong, X. (2022) Improved pea reference genome and pan-genome highlight genomic features and evolutionary characteristics. Nat. Genet. 54, 1553–1563.

Ye, Q., Zhu, F., Sun, F., Wang, T.C., Wu, J., Liu, P., Shen, C., Dong, J. and Wang, T. (2022) Differentiation trajectories and biofunctions of symbiotic and un-symbiotic fate cells in root nodules of *Medicago truncatula*. Mol. Plant 15, 1852–1867.

Zhang, L., He, C., Lai, Y., Wang, Y., Kang, L., Liu, A., Lan, C., Su, H., Gao, Y., Li, Z., Yang, F., Li, Q., Mao, H., Chen, D., Chen, W., Kaufmann, K. and Yan, W. (2023) Asymmetric gene expression and cell-type-specific regulatory networks in the root of bread wheat revealed by single-cell multiomics analysis. Genome Biol. 24, 65.

Zhang, T.Q., Chen, Y., Liu, Y., Lin, W.H. and Wang, J.W. (2021a) Single-cell transcriptome atlas and chromatin accessibility landscape reveal differentiation trajectories in the rice root. Nat. Commun. 12, 2053.

Zhang, T.Q., Chen, Y. and Wang, J.W. (2021b) A single-cell analysis of the *Arabidopsis* vegetative shoot apex. Dev. Cell 56, 1–19.

Zhang, T.Q., Lian, H., Zhou, C.M., Xu, L., Jiao, Y. and Wang, J.W. (2017) A two-step model for de novo activation of *WUSCHEL* during plant shoot regeneration. Plant Cell 29, 1073–1087.

