## Supplemental Figures for "Single-cell transcriptomic analysis of pea shoot development and cell-type-specific responses to boron deficiency"

### Supplementary Figures

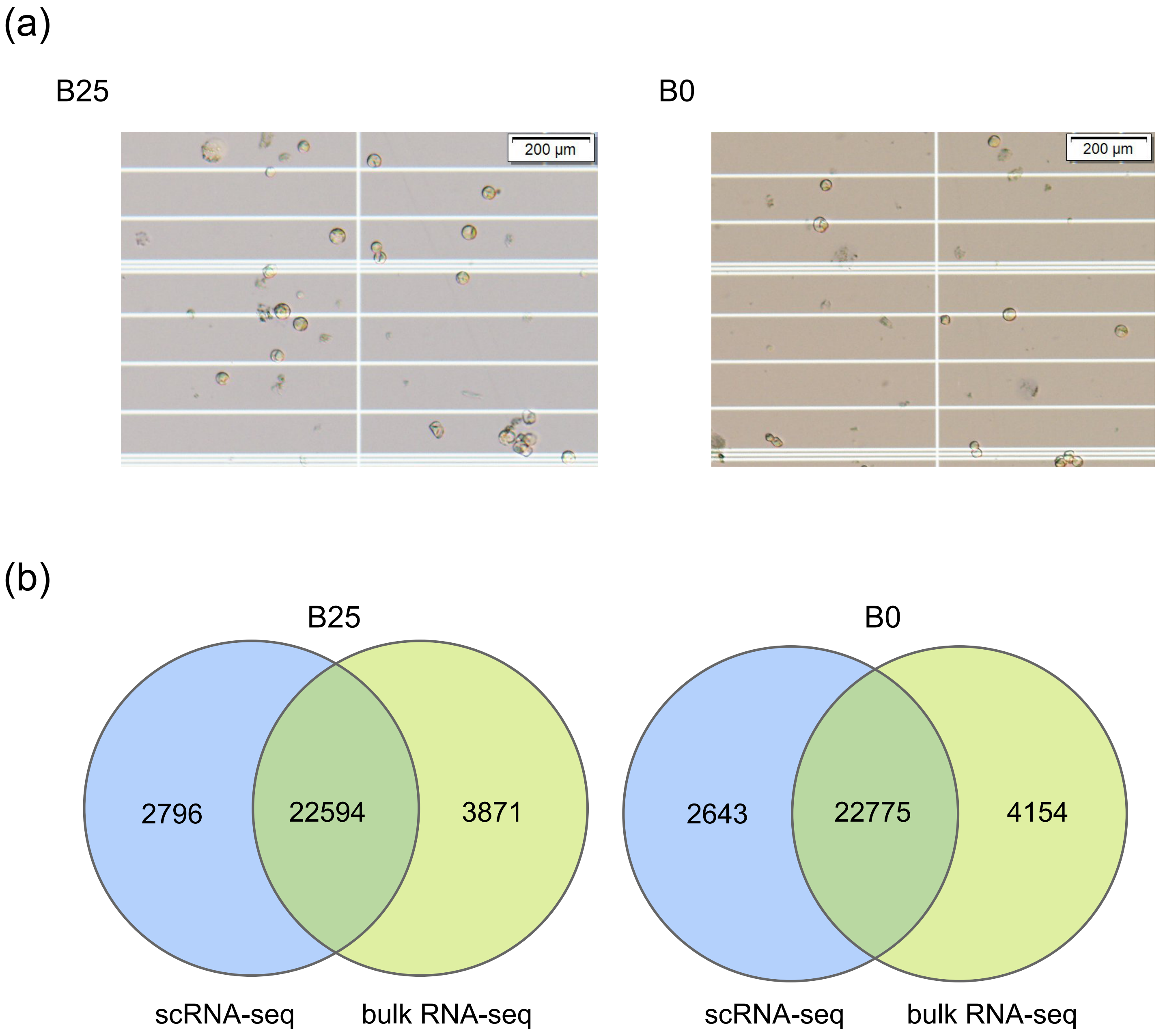


**Figure S1** Overview of scRNA-seq analysis. (a) Protoplast preparations for scRNA-seq. (b) Venn diagram showing numbers of unique and common genes detected in bulk RNA-seq and scRNA-seq in B0 and B25 samples.

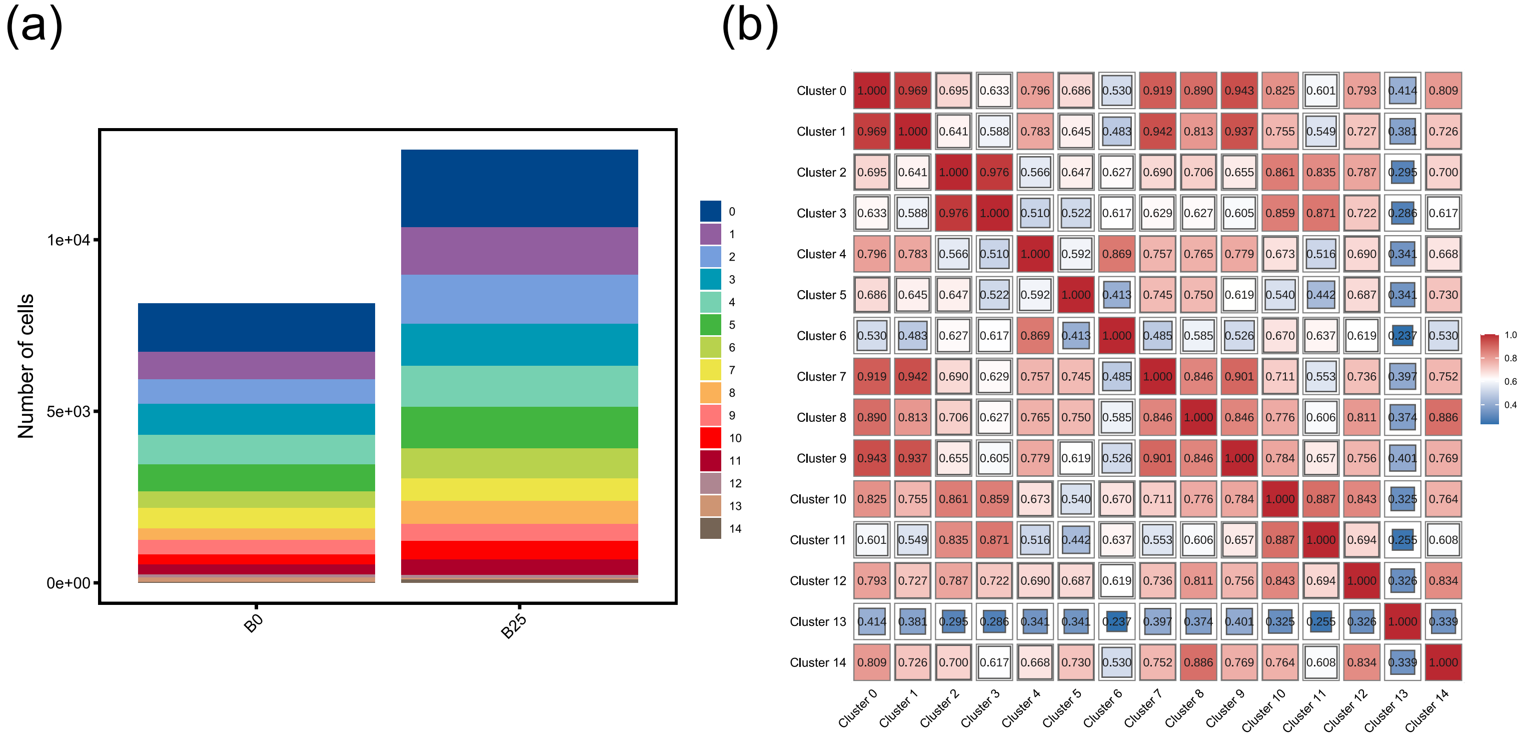


**Figure S2** Numbers of cells in each scRNA-seq cluster and correlation of each cluster. (a) Number of cells in each cluster in B0 and B25 samples. (b) Correlations between 15 cell clusters expressed as proportion of detected genes in common (greater than 0.5, red; less than 0.5, blue).

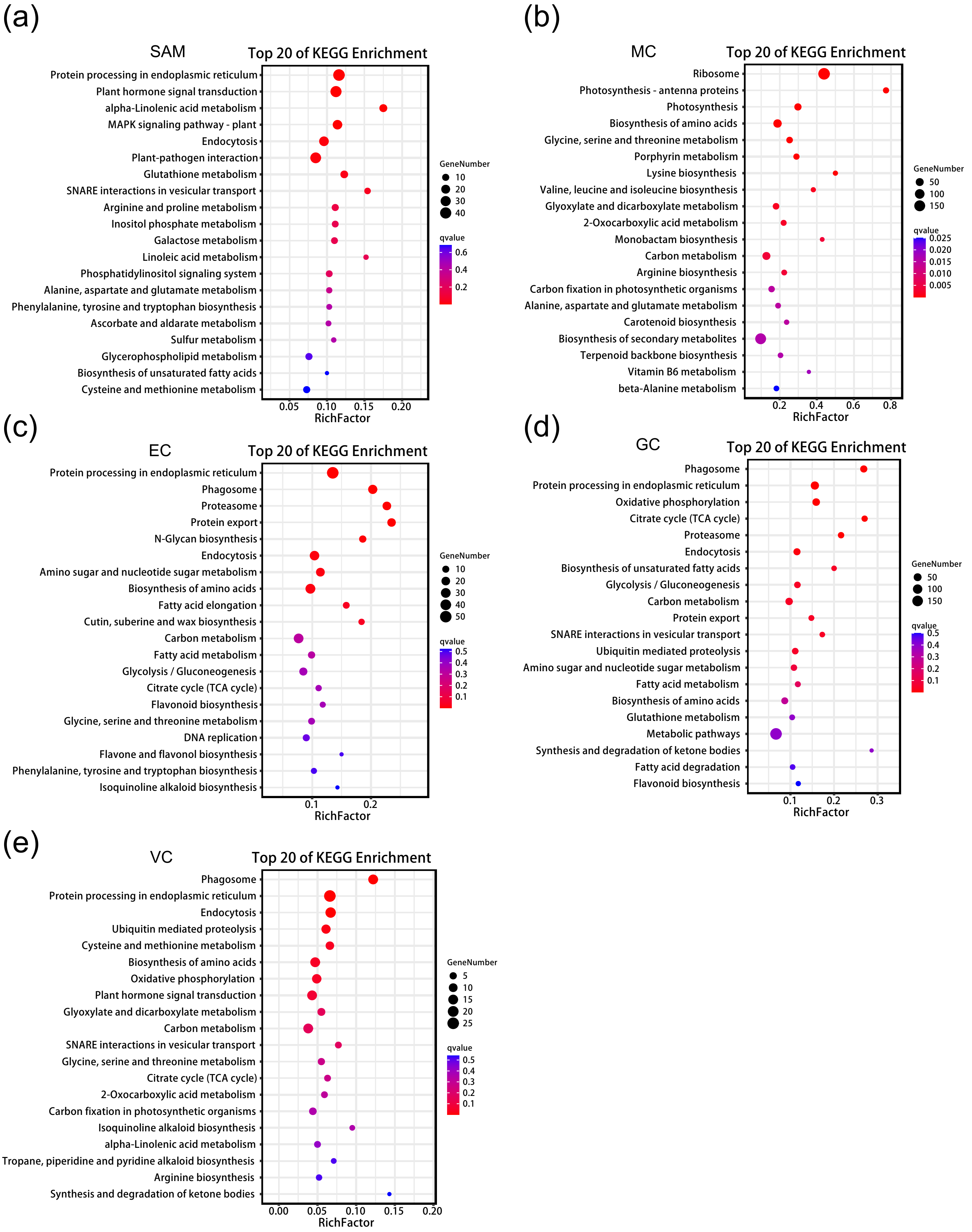


**Figure S3** KEGG enrichment annotation of genes expressed specifically in the five known cell types. (a) SAM (b) MC (c) EC (d) GC (e) VC.

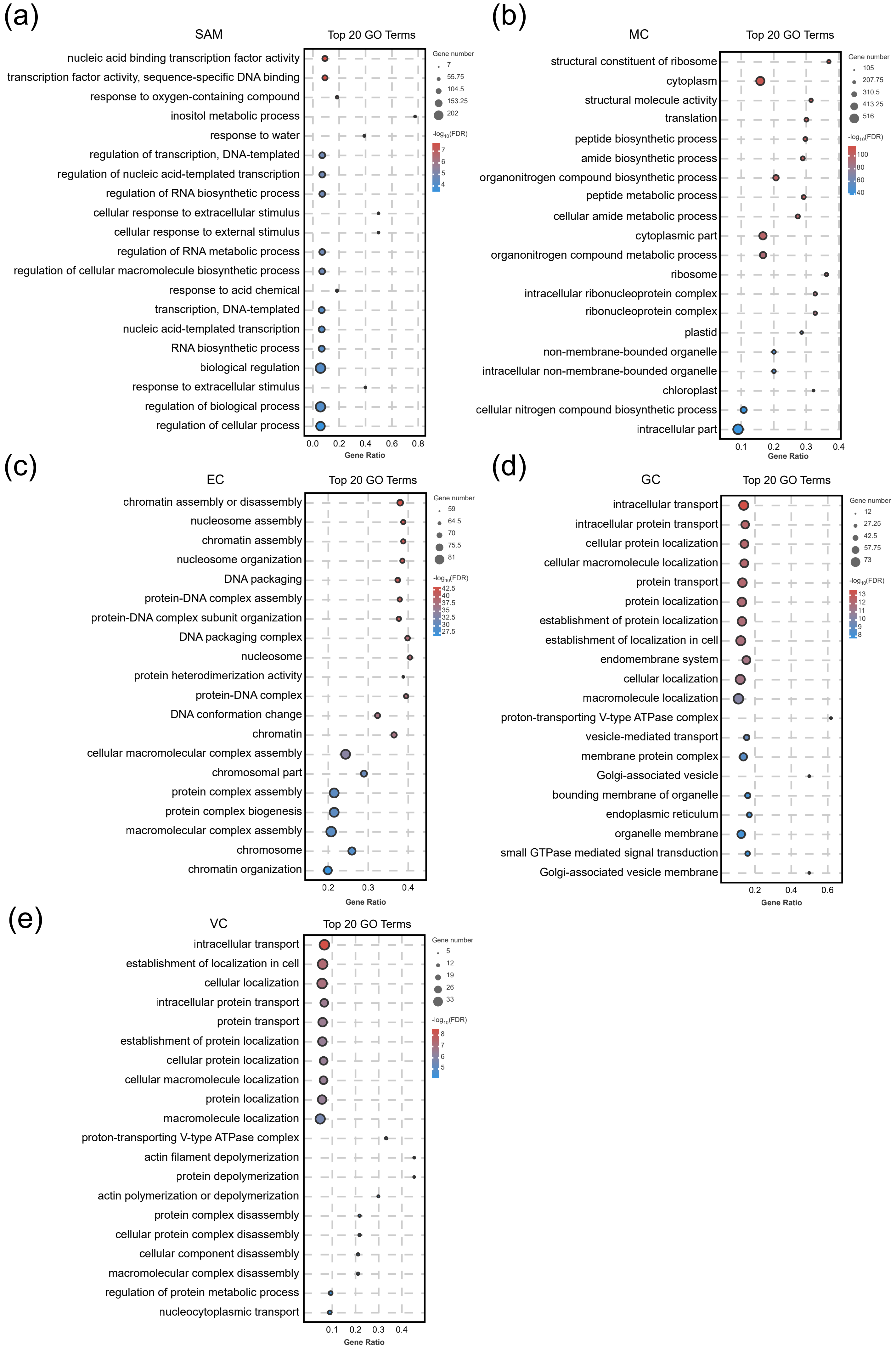


**Figure S4** GO term enrichment annotation of genes expressed specifically in the five known cell types. (a) SAM (b)MC (c) EC (d) GC (e) VC.

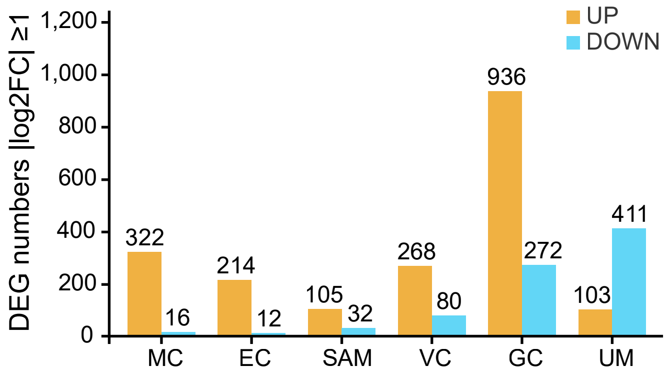


**Figure S5** Numbers of DEGs in response to B-deficiency in six cell types. DEGs were identified as up (orange) or down (blue) in B0 relative to B25 with a fold-change of $|log_2FC| \geq 1$.

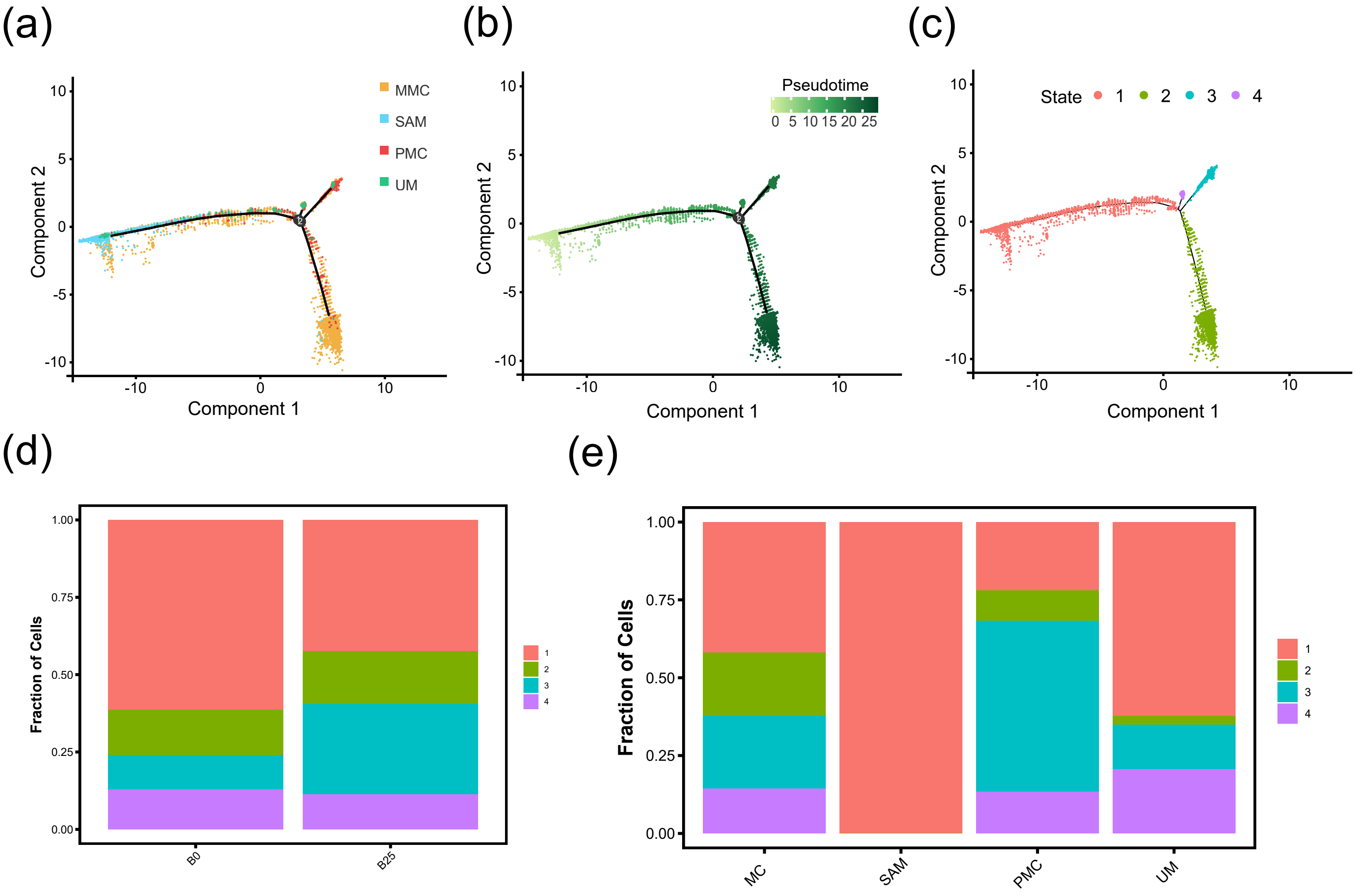


**Figure S6** Pseudotime trajectory from SAM to MC. (a-c) The distribution of cells along the pseudotime trajectory color-coded by cell type (a), pseudotime states, represented by intensity of green (b), and branch states, color-coded (c). MMC, mature mesophyll cells; SAM, shoot apical meristem cells; PMC, proliferating mesophyll cells; UM, unknown meristem. (d-e) Cell proportion of four states in B0 and B25 samples (d), and four different cell types (e).

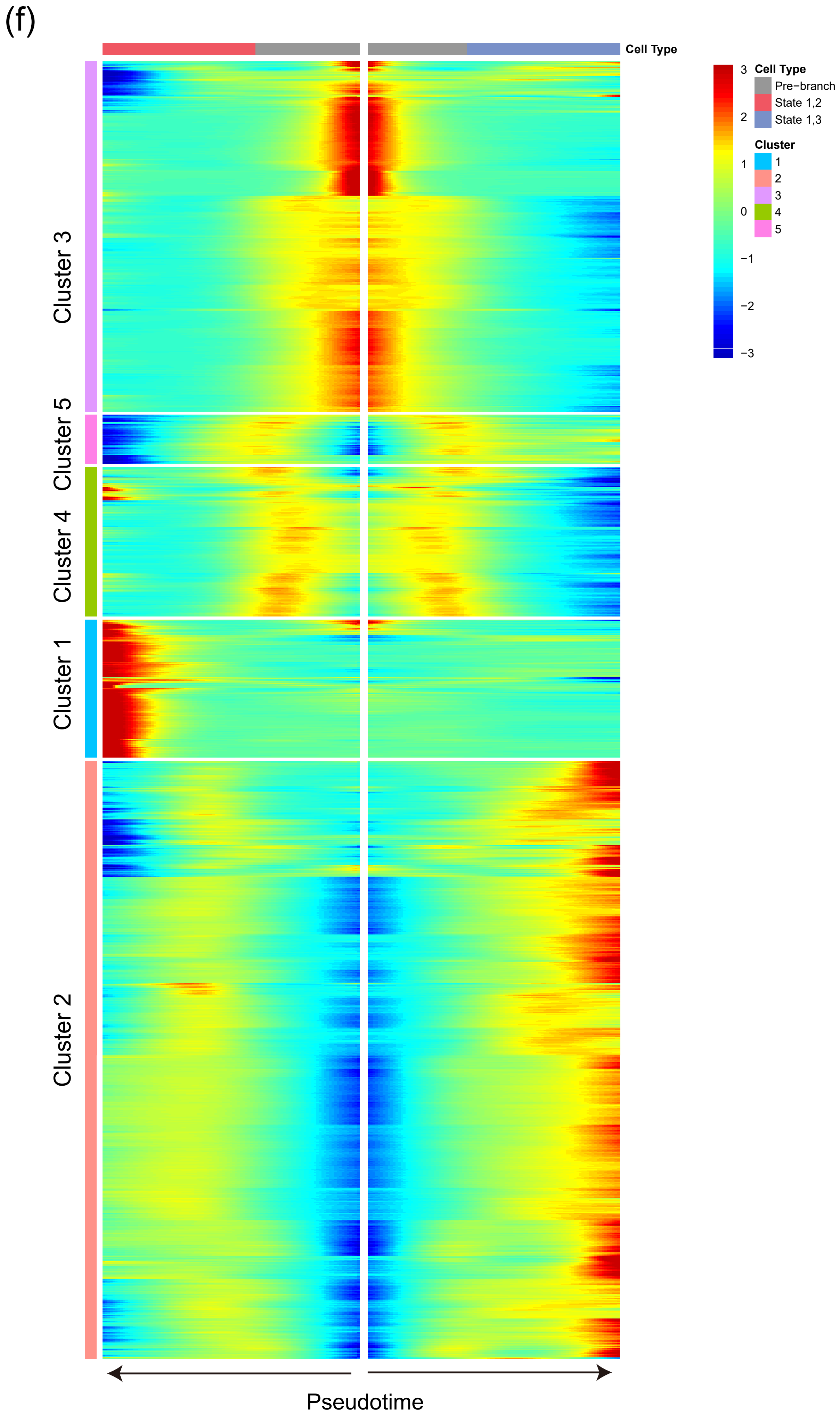


**Figure S6** (f) Heatmap showing all gene clusters over the pseudotime trajectories.
